# EvoRx: Machine Learning-Based Prediction of Antibiotic Resistance in Escherichia coli Using Genomic Features

**DOI:** 10.64898/2026.11.16.767019

**Authors:** Aarna Arora, Elamathi Natarajan

## Abstract

Antimicrobial resistance (AMR) complicates bacterial infection management, motivating genomic approaches to predicting antibiotic susceptibility. This study assessed the predictive performance and generalisability of machine-learning models using resistance-associated genomic features from Escherichia coli. Publicly available genomic and phenotypic records yielded 446 genomes, 252 binary genomic features and 1,935 susceptibility observations across nine antibiotics. Logistic Regression, Random Forest and XGBoost classifiers were compared using validation ROC-AUC, followed by evaluation on held-out test subsets. Additional analyses examined identical genomic feature profiles across partitions, performance across source publications and feature redundancy. Ampicillin and trimethoprim-sulfamethoxazole achieved held-out accuracies of 93.98% and 93.59%, ROC-AUC values of 0.912 and 0.954, and resistant-isolate recall of 95.16% and 94.55%, respectively. In contrast, ceftazidime achieved 88.52% accuracy and 0.940 ROC-AUC but identified only 22.22% of resistant isolates. Seventy-four held-out observations shared identical 252-feature profiles with training isolates; all were correctly classified. After excluding overlapping observations without retraining, ROC-AUC decreased from 0.912 to 0.848 for ampicillin and from 0.954 to 0.901 for trimethoprim-sulfamethoxazole. Exploratory cross-publication evaluations showed variable performance, with shared multilocus sequence types in 16 of 18 comparisons. Reducing genomic features did not materially alter validation performance in the two antibiotics examined. These findings indicate that genomic predictors can discriminate resistance phenotypes, but aggregate accuracy may obscure poor resistant-isolate detection. The findings also distinguish performance on conventionally held-out isolates from performance on previously unseen resistance-feature profiles, an important consideration when bacterial population structure and correlated determinants substantially influence model evaluation. Independent phylogenetic and prospective validation remains necessary before clinical use.

## 1. Introduction

### 1.1 The Global Challenge of Antimicrobial Resistance

Antimicrobial resistance (AMR) compromises the treatment of bacterial infections by reducing the effectiveness of existing antibiotics. Resistance arises through genetic mutations or the acquisition of resistance determinants, with antimicrobial exposure selecting for organisms capable of surviving treatment. The clinical consequences include prolonged infections, treatment failure, increased healthcare expenditure and mortality. In 2019, bacterial AMR was estimated to be directly responsible for 1.27 million deaths worldwide and associated with approximately 4.95 million deaths [1]. Global projections indicate that the burden will remain substantial through 2050 [2]. Improving the detection and surveillance of resistant pathogens is therefore a priority, particularly as advances in bacterial genomics offer opportunities to complement conventional susceptibility testing with computational prediction.

### 1.2 Antibiotic Resistance in Escherichia coli

Escherichia coli is a Gram-negative bacterium that commonly inhabits the intestinal tract but also causes urinary tract infections, bloodstream infections and other invasive diseases. It is among the leading bacterial pathogens associated with global mortality [3]. Resistance in E. coli spans several antibiotic classes, including penicillins, cephalosporins, fluoroquinolones and folate-pathway inhibitors. Its extensive genomic diversity and capacity for horizontal gene transfer facilitate the acquisition and dissemination of resistance determinants. However, the distribution of these determinants differs between bacterial lineages, and genetically similar isolates may exhibit different susceptibility phenotypes. The availability of publicly accessible genomic sequences and associated antimicrobial susceptibility data provides a basis for investigating these relationships computationally.

### 1.3 Genomic Mechanisms Underlying Antibiotic Resistance

Antibiotic resistance in E. coli involves several mechanisms, principally enzymatic inactivation, modification of antibiotic targets, reduced membrane permeability and active efflux. Beta-lactam resistance frequently involves beta-lactamases encoded by genes such as blaTEM and blaCTX-M, although the antibiotics affected depend on the specific enzyme variant [4]. Resistance to trimethoprim-sulfamethoxazole may involve dfr and sul genes, which encode altered enzymes in the folate biosynthesis pathway. Chromosomal mutations, regulatory variation and interactions among resistance determinants can further modify susceptibility.

Genomic resources including CARD, ResFinder and AMRFinderPlus enable the identification of established resistance-associated genes and mutations [5–7]. Hunt et al. developed ARIBA, a computational method for identifying resistance determinants directly from sequencing reads [8]. Nevertheless, the presence of a resistance-associated sequence does not invariably correspond to phenotypic resistance. Gene expression, sequence functionality and the broader genomic background may influence whether a detected determinant produces a measurable resistance phenotype.

### 1.4 Current Methods of Antimicrobial Susceptibility Testing

Phenotypic antimicrobial susceptibility testing (AST) remains the reference approach for determining bacterial responses to antibiotic exposure. Common methods include disk diffusion, broth microdilution and automated susceptibility testing systems. Disk diffusion measures inhibition-zone diameters, whereas broth microdilution determines the minimum inhibitory concentration (MIC). Results are interpreted using established breakpoints, including those published by EUCAST and CLSI.

Although phenotypic AST directly measures susceptibility, it generally requires bacterial cultivation and laboratory processing. Whole-genome sequencing provides an alternative source of information by identifying genetic determinants associated with resistance. However, incomplete knowledge of resistance mechanisms, variation in gene expression and limitations in genomic annotation can produce discrepancies between predicted and observed phenotypes [9]. Genomic approaches therefore require validation against laboratory susceptibility measurements rather than being treated as equivalent to phenotypic AST.

### 1.5 Machine Learning in Genomic AMR Prediction

Supervised machine learning (ML) has been investigated as a means of predicting antimicrobial resistance from bacterial genomic characteristics. By analysing genomic features alongside known susceptibility phenotypes, classification algorithms can identify statistical relationships that extend beyond individual resistance-gene associations. Moradigaravand et al. demonstrated the feasibility of resistance prediction in E. coli using large-scale pan-genome data [10]. Pearcy et al. subsequently combined ML with genome-scale metabolic modelling to investigate genetic determinants and metabolic adaptations associated with resistance [11].

Logistic Regression, Random Forest and gradient-boosted decision trees offer different approaches to modelling these relationships. However, predictive performance depends on the composition of the training data, resistance prevalence, feature representation and validation strategy. Kim et al. examined these limitations in genomic AMR prediction, including challenges involving model interpretability and clinical applicability [12]. Bacterial population structure presents an additional concern: related isolates may share genomic characteristics across training and evaluation datasets, allowing models to exploit lineage-associated patterns rather than associations that generalise to genetically distinct populations [13].

### 1.6 Research Gaps, Study Objectives and Hypotheses

Existing genomic ML approaches have demonstrated predictive utility, but several limitations warrant further investigation. Overall accuracy may obscure poor identification of resistant isolates, particularly in datasets with unequal class distributions. Performance may also differ substantially between antibiotics because of differences in resistance mechanisms and available phenotypic observations. Furthermore, identical genomic feature profiles across training and test datasets can complicate the interpretation of model generalisability. These considerations motivate antibiotic-specific evaluation alongside analyses of genomic profile overlap and population structure.

#### Research Question

To what extent can resistance-associated genomic features predict antibiotic resistance phenotypes in E. coli across nine antibiotics, and how reliably do these predictions generalise to isolates with previously unseen genomic feature profiles?

#### Research Objectives

1. To construct a genomic dataset comprising 446 E. coli genomes, 252 binary resistance-associated features and corresponding antibiotic susceptibility phenotypes.
2. To develop and compare Logistic Regression, Random Forest and XGBoost models for resistance classification across nine antibiotics.
3. To evaluate antibiotic-specific predictive performance using accuracy, ROC-AUC and resistant-isolate recall.
4. To assess the influence of overlapping training-test genomic feature profiles and investigate generalisation through exploratory cross-publication validation.
5. To examine associations between genomic resistance determinants and phenotypic resistance, alongside the effects of feature redundancy on model performance.

#### Hypotheses

##### Primary Hypothesis (H1)

Resistance-associated genomic features provide predictive information for distinguishing resistant and susceptible E. coli isolates, with discriminatory performance varying across antibiotics.

##### Generalisation Hypothesis (H2)

Predictive performance differs when test isolates sharing identical genomic feature profiles with training isolates are excluded, indicating that feature-profile overlap affects estimates of model generalisability.

##### Null Hypothesis (H0)

Resistance-associated genomic features do not provide predictive discrimination beyond chance for the antibiotic susceptibility phenotypes examined.

## 2. Materials and Methods

### 2.1 Study Design and Computational Workflow

EvoRx was designed as a retrospective computational study examining the prediction of phenotypic antibiotic resistance in Escherichia coli from genomic resistance determinants. The study combined publicly available bacterial genome assemblies with experimentally derived antimicrobial susceptibility records to develop antibiotic-specific binary classification models.

The computational workflow comprised five stages: acquisition and quality assessment of genomic and phenotypic records; identification of resistance-associated genes and mutations; construction of antibiotic-specific feature matrices; supervised model development and evaluation; and supplementary analyses of genomic profile overlap, cross-publication performance and feature redundancy. Logistic Regression, Random Forest and XGBoost were compared using a common genomic feature representation. Predictive performance was assessed separately for nine antibiotics to account for differences in phenotype availability, resistance prevalence and underlying genetic mechanisms.

### 2.2 Dataset Acquisition and Construction

Antimicrobial susceptibility records were retrieved from the Bacterial and Viral Bioinformatics Resource Centre (BV-BRC), formerly PATRIC, through its genome_amr application programming interface. Records were queried for E. coli (taxonomic identifier 562) across an initial panel of 24 antibiotics. Only observations identified as originating from laboratory methods were retained; computationally inferred susceptibility records were excluded.

Genome metadata were obtained from the BV-BRC genome database. Eligible assemblies required an identifiable NCBI accession, a BV-BRC genome-quality designation of Good, CheckM-estimated completeness of at least 95%, contamination not exceeding 5%, and no deprecated status. After phenotype cleaning and restriction to the quality-filtered genome collection, 456 genomes possessed at least one usable susceptibility observation.

Genomes were prioritised according to the number of antibiotics with available phenotypic measurements. Corresponding nucleotide assemblies were downloaded from NCBI using the Datasets command-line interface. Ten genomes were subsequently excluded because AMRFinderPlus screening did not complete within the specified 900-second execution limit, including retry attempts. The final dataset comprised 446 successfully screened genomes with unique assembly accessions.

### 2.3 Antibiotic Susceptibility Data and Phenotype Labelling

Susceptibility phenotypes were obtained from experimentally derived BV-BRC records and standardised into two categories: resistant (R) and susceptible (S). Intermediate and nonsusceptible classifications were retained separately during data processing but excluded from the binary modelling dataset. Records without interpretable phenotypes were also excluded.

Repeated genome-antibiotic observations were consolidated during phenotype processing, with conflicting classifications assessed before dataset construction. The final dataset contained 1,935 unambiguous genome-antibiotic phenotype observations, with no conflicting duplicate classifications remaining.

Nine antibiotics were represented: amoxicillin-clavulanic acid, ampicillin, cefazolin, cefotaxime, cefoxitin, ceftazidime, ceftriaxone, piperacillin-tazobactam and trimethoprim-sulfamethoxazole. Susceptibility coverage differed across antibiotics, ranging from 37 labelled genomes for cefoxitin to 416 for ampicillin.

The source records included susceptibility measurements obtained under different laboratory methods and interpretive standards, including CLSI and EUCAST. Because breakpoint definitions and testing procedures were not harmonised retrospectively, the reported laboratory classifications were used as the reference phenotypes.

### 2.4 Genomic Feature Extraction

Resistance-associated genomic features were identified using NCBI AMRFinderPlus version 4.2.7, with database release 2026-08-07.1. Genome assemblies were analysed in nucleotide mode using the Escherichia-specific configuration and the--plus option. This procedure enabled identification of curated antimicrobial resistance genes and supported point mutations.

AMRFinderPlus outputs were parsed into a genome-by-feature matrix. Each detected resistance determinant was encoded as 1 and its absence from a successfully screened assembly as 0. Failed genomic screens were not interpreted as negative detections; the corresponding genomes were excluded from the final cohort.

The resulting matrix contained 252 binary resistance-associated features across 446 genomes. Stress-response and virulence annotations were maintained separately rather than incorporated into the resistance-feature matrix. Genomic accession identifiers were retained for record linkage but excluded from model predictors.

### 2.5 Data Cleaning, Quality Control and Preprocessing

Genomic features, phenotype records and isolate metadata were integrated using NCBI assembly accessions. The final dataset was checked for unique accession identifiers, consistent feature encoding and correspondence between genomic and phenotypic records.

Genome-quality filtering was performed before feature extraction using BV-BRC quality annotations and CheckM estimates. Only assemblies with successful AMRFinderPlus screening were retained, preventing incomplete annotation runs from being misrepresented as genomes lacking resistance determinants.

Phenotype records without definitive resistant or susceptible classifications were omitted rather than imputed. For each antibiotic, isolates lacking a corresponding phenotype were excluded from that antibiotic’s modelling dataset while remaining eligible for analyses of other antibiotics.

The 252 genomic features were retained as binary predictors. Isolate identifiers, publication information, multilocus sequence types and laboratory testing metadata were excluded from the predictor matrices. These variables were preserved separately for dataset partitioning and subsequent validation analyses.

### 2.6 Training, Validation and Test Partitioning

Separate training, validation and test partitions were generated for each antibiotic using stratified group-based splitting implemented through scikit-learn’s StratifiedGroupKFold.

An outer five-fold procedure, initialised with random seed 42, designated one fold as the held-out test subset. The remaining observations underwent an inner four-fold split using random seed 43, assigning one fold to validation and the remainder to training. This produced approximate training, validation and test proportions of 60%, 20% and 20%, respectively, although exact proportions differed between antibiotics because group boundaries were preserved.

Grouping was based on multilocus sequence type (MLST) when available; otherwise, the BioSample-derived isolate-group identifier was used. This prevented isolates sharing the recorded grouping identifier from being distributed across different partitions.

Partition assignments were recorded in a split manifest containing isolate accessions, phenotype labels, MLST information, source-publication identifiers and group assignments. The test subsets were reserved for evaluation after model selection.

Although the partitioning procedure reduced overlap between recorded groups, MLST-based separation did not establish whole-genome phylogenetic independence. Accordingly, additional analyses examined identical genomic feature profiles across training and test partitions and assessed performance across publication-defined subsets.

### 2.7 Machine-Learning Algorithms

Three supervised classification algorithms were evaluated: Logistic Regression (LR), Random Forest (RF) and Extreme Gradient Boosting (XGBoost). Each algorithm was trained separately for the nine antibiotic susceptibility outcomes using the same 252 binary genomic features.

Logistic Regression modelled resistance probability through a linear combination of genomic predictors. Random Forest combined multiple decision trees to accommodate nonlinear relationships and interactions between resistance determinants. XGBoost employed sequentially constructed, gradient-boosted decision trees to minimise classification error through an iterative optimisation procedure.

The resistant phenotype was designated the positive classification outcome. Model development was conducted independently for each antibiotic, allowing algorithm selection to reflect differences in phenotype distributions and genomic associations.

### 2.8 Model Selection and Hyperparameter Optimisation

Candidate algorithms were compared using their performance on the designated validation subset. The principal selection criterion was the area under the receiver operating characteristic curve (ROC-AUC), which measures discrimination between resistant and susceptible isolates across classification thresholds.

For each antibiotic, the algorithm achieving the highest validation ROC-AUC was selected for final evaluation. The selected model was subsequently applied to the held-out test subset, which was not used to determine the preferred algorithm.

The archived experiments document validation-based algorithm selection rather than a systematic hyperparameter-search procedure. Accordingly, no independent hyperparameter optimisation is attributed to the reported comparisons.

### 2.9 Performance Evaluation Metrics

Predictive performance was assessed using accuracy, balanced accuracy, precision, recall, F1-score and ROC-AUC. Resistance recall, equivalent to sensitivity for the resistant class, measured the proportion of phenotypically resistant isolates correctly identified. Balanced accuracy was calculated as the mean of resistant-class sensitivity and susceptible-class specificity.

ROC-AUC quantified the ability of predicted scores to discriminate between resistant and susceptible isolates independently of a single classification threshold. Confusion-matrix-derived metrics were used to distinguish overall classification performance from resistance detection.

Uncertainty in selected test metrics was assessed using bootstrap-derived 95% confidence intervals. Results were reported separately for each antibiotic because sample sizes and class distributions differed substantially.

### 2.10 Genomic Profile Overlap and Leakage Sensitivity Analyses

Potential overlap between training and test data was examined at both the metadata and genomic feature levels. Recorded MLST, BioSample accession and isolate-group identifiers were compared across partitions to identify shared grouping information.

A separate genomic profile audit compared the complete 252-feature binary vectors of training and test isolates. Test isolates whose feature vectors exactly matched at least one training isolate were identified, irrespective of their recorded isolate identifiers.

To assess the sensitivity of reported performance to this overlap, test isolates with matching training profiles were excluded, and the previously fitted models were evaluated on the remaining test observations. Models were not retrained during this analysis.

Identical resistance-feature profiles were distinguished from whole-genome identity, since the selected feature representation did not capture all genomic variation.

### 2.11 Exploratory Cross-Publication Validation

An additional evaluation examined model performance across publication-defined isolate groups. For each eligible antibiotic and source publication, isolates associated with that publication were designated as the evaluation subset, while isolates from other publications formed the training population.

Publication-held-out experiments were conducted only where the available observations supported meaningful binary classification assessment. Performance was summarised using accuracy, balanced accuracy, resistant recall and ROC-AUC.

The analysis comprised 18 evaluable publication-held-out comparisons across seven antibiotics. Shared MLST sequence types between training and evaluation publications were assessed to characterise residual population overlap.

Because model-selection procedures were not fully nested within publication-held-out evaluation, these experiments were interpreted as exploratory assessments of cross-publication performance rather than independent external validation.

### 2.12 Genomic Feature Interpretation and Redundancy Analysis

Genomic feature interpretation examined associations between individual resistance determinants and observed susceptibility phenotypes. For ampicillin, gene prevalence was compared between resistant and susceptible isolates to identify determinants showing differential representation.

Feature redundancy was investigated by comparing the original 252-feature representation with reduced feature sets for ampicillin and trimethoprim-sulfamethoxazole. The reduced matrices contained 144 and 138 features, respectively.

Predictive performance was compared using validation accuracy, balanced accuracy and ROC-AUC. These analyses assessed the sensitivity of model performance to redundant genomic information, without treating predictive associations as evidence of causal resistance mechanisms.

### 2.13 Software, Computational Environment and Reproducibility

Machine-learning analyses were performed in Python 3.13.16 using scikit-learn 1.6.1, XGBoost 3.4.1, pandas 2.2.3 and NumPy 2.1.3.

The research archive retained antibiotic-specific modelling datasets, a deterministic partition manifest, validation results, selected model objects, feature-order records, held-out test metrics and supplementary validation outputs. Trained models were preserved in Joblib format.

Additional archived materials included bootstrap confidence intervals, genomic profile-overlap audits, publication-held-out evaluation results and feature-redundancy analyses. These records provided the basis for examining model performance and the sensitivity of reported results to dataset structure.

## 3. Results

### 3.1 Final Dataset Characteristics and Antibiotic Coverage

The final EvoRx dataset comprised 446 unique Escherichia coli genomes, each represented by 252 binary resistance-associated genomic features. Across the nine antibiotics, 1,935 genome-antibiotic observations had definitive resistant or susceptible phenotype labels. Antibiotic coverage was unequal, ranging from 37 observations for cefoxitin to 416 for ampicillin.

Ampicillin had the largest modelling dataset (n = 416) followed by trimethoprim-sulfamethoxazole (n = 386), ceftriaxone (n = 337) and ceftazidime (n = 305) Smaller datasets were available for cefazolin (n = 135), piperacillin-tazobactam (n = 130) amoxicillin-clavulanic acid (n = 105), cefotaxime (n = 84) and cefoxitin (n = 37)

Phenotype distributions also varied. Ampicillin and trimethoprim-sulfamethoxazole datasets contained predominantly resistant isolates, whereas susceptible isolates predominated for ceftazidime, ceftriaxone and piperacillin-tazobactam. These differences were retained in the antibiotic-specific training, validation and test partitions.

### 3.2 Comparison of Machine-Learning Algorithms

Logistic Regression, Random Forest and XGBoost were compared independently for each antibiotic using validation ROC-AUC. Logistic Regression was selected for five antibiotics: ampicillin, cefazolin, cefotaxime, piperacillin-tazobactam and trimethoprim-sulfamethoxazole. Random Forest was selected for amoxicillin-clavulanic acid, cefoxitin, ceftazidime and ceftriaxone. XGBoost was not selected as the final model for any antibiotic.

For ampicillin, Logistic Regression achieved a validation ROC-AUC of 0.901, compared with 0.877 for Random Forest and 0.823 for XGBoost. Logistic Regression also produced the highest validation ROC-AUC for cefazolin (0.858) and trimethoprim-sulfamethoxazole (0.878).

Random Forest achieved the highest validation ROC-AUC for ceftazidime (0.856) and ceftriaxone (0.866). For amoxicillin-clavulanic acid, its validation ROC-AUC was 0.764, compared with 0.707 for Logistic Regression and 0.659 for XGBoost.

Validation discrimination was less consistent for the smaller datasets. The selected cefotaxime model had a validation ROC-AUC of 0.431, while the selected cefoxitin model achieved 0.500. These values differed considerably from the subsequent held-out test results.

### 3.3 Held-Out Test Performance Across Nine Antibiotics

The selected models were evaluated on antibiotic-specific held-out test subsets containing between 9 and 83 isolates. Test accuracy ranged from 67.86% for cefazolin to 100% for cefoxitin, while ROC-AUC ranged from 0.795 to 1.000.

**Table 1.** Held-out test performance of the selected EvoRx models.

| **Antibiotic** | **Selected Model** | **Test n** | **Accuracy (%)** | **ROC-AUC** | **Resistant recall** |
|---|---|---|---|---|---|
| Amoxicillin-clavulanic acid | RF | 22 | 86.36 | 0.915 | 0.929 |
| Ampicillin | LR | 83 | 93.98 | 0.912 | 0.952 |
| Cefazolin | LR | 28 | 67.86 | 0.795 | 0.533 |
| Cefotaxime | LR | 20 | 85.00 | 1.000 | 0.700 |
| Cefoxitin | RF | 9 | 100.00 | 1.000 | 1.000 |
| Ceftazidime | RF | 61 | 88.52 | 0.940 | 0.222 |
| Ceftriaxone | RF | 74 | 83.78 | 0.889 | 0.500 |
| Piperacillin-tazobactam | LR | 26 | 92.31 | 1.000 | 0.500 |
| Trimethoprim-sulfamethoxazole | LR | 78 | 93.59 | 0.954 | 0.945 |

*LR, Logistic Regression; RF, Random Forest. Resistant recall represents sensitivity for the resistant phenotype.*

**Figure 1.**
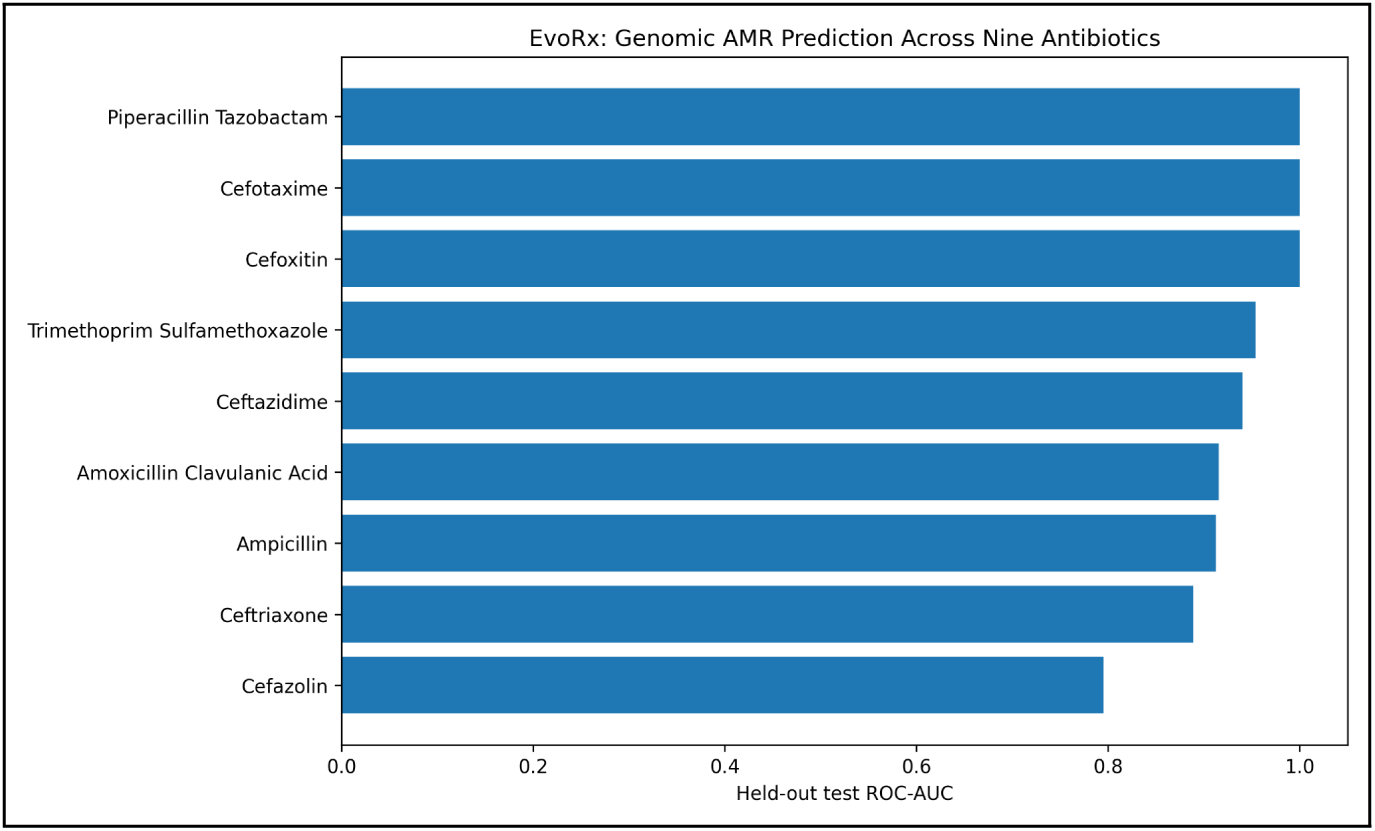
Held-out test ROC-AUC of the selected EvoRx classification models across nine antibiotics. Models were selected using validation ROC-AUC and evaluated on antibiotic-specific held-out test subsets. Test sample sizes ranged from 9 to 83 isolates.

Ampicillin and trimethoprim sulfamethoxazole produced accuracies above 93%, with ROC-AUC values exceeding 0.91. Amoxicillin-clavulanic acid achieved 86.36% accuracy and a ROC-AUC of 0.915. Cefazolin had the lowest overall accuracy (67.86%) and ROC-AUC (0.795).

Three antibiotics-cefotaxime, cefoxitin and piperacillin-tazobactam achieved a test ROC-AUC of 1.000. Their test subsets contained 20, 9 and 26 isolates, respectively.

### 3.4 Resistance Detection and Class-Specific Performance

Resistant-isolate recall varied more widely than overall accuracy, ranging from 0.222 for ceftazidime to 1.000 for cefoxitin.

**Figure 2.**
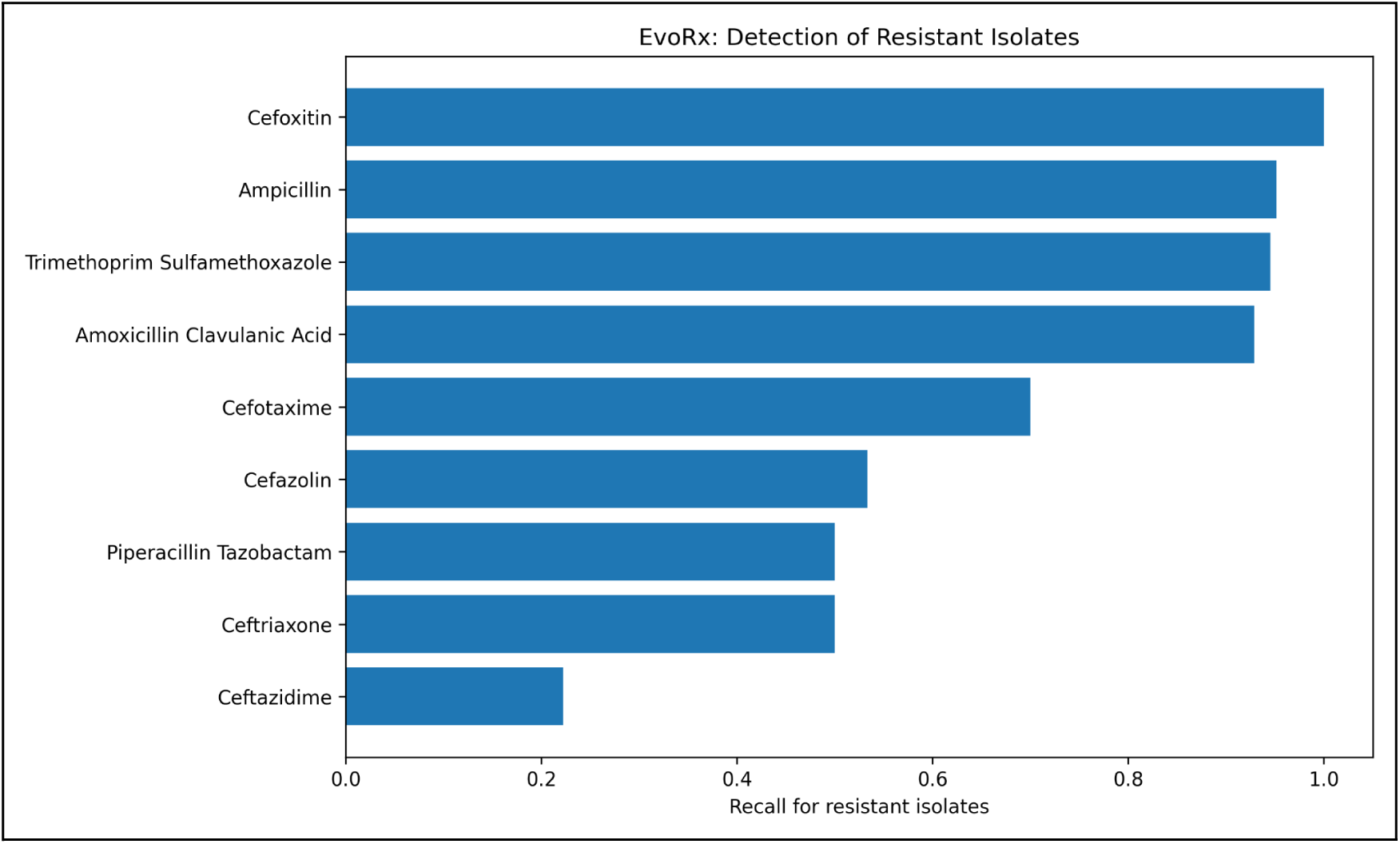
Resistant-isolate recall of the selected EvoRx models across nine antibiotics. Recall represents the proportion of phenotypically resistant isolates correctly classified as resistant in the held-out test subsets.

The ampicillin model correctly classified 59 of 62 resistant isolates and 19 of 21 susceptible isolates, producing resistant recall of 0.952 and balanced accuracy of 0.928. For trimethoprim-sulfamethoxazole, resistant recall was 0.945 and balanced accuracy was 0.929.

Ceftazidime showed the largest discrepancy between overall accuracy and resistance detection. Although accuracy reached 88.52% and ROC-AUC was 0.940, only 2 of 9 resistant isolates were correctly classified. All 52 susceptible isolates were correctly identified, giving a balanced accuracy of 0.611.

For ceftriaxone, the model correctly identified 12 of 24 resistant isolates and all 50 susceptible isolates, resulting in resistant recall of 0.500 and balanced accuracy of 0.750. The piperacillin-tazobactam model identified 2 of 4 resistant isolates, despite achieving 92.31% overall accuracy.

Cefazolin produced a recall of 0.533, with 8 of 15 resistant isolates correctly classified. The cefotaxime model identified 7 of 10 resistant isolates, while the cefoxitin model correctly classified all nine test observations, including seven resistant isolates.

### 3.5 Impact of Overlapping Genomic Feature Profiles

An audit of the 252-feature genomic representation identified 74 held-out test observations whose feature profiles were identical to those of at least one training isolate. These observations occurred across six antibiotic-specific datasets.

The largest number of matching profiles occurred for trimethoprim sulfamethoxazole (27 of 78 test observations), followed by ceftazidime (18 of 61), ampicillin (15 of 83), ceftriaxone (9 of 74), piperacillin-tazobactam (4 of 26) and cefoxitin (1 of 9). No matching training profiles were detected for amoxicillin-clavulanic acid, cefazolin or cefotaxime.

All 74 overlapping test observations were correctly classified in the original held-out evaluation. The overlap audit identified 57 observations whose matching training profiles were associated with the same source publication. No matching MLST, BioSample accession or isolate-group identifiers were identified among the recorded metadata comparisons.

The profile audit also identified two trimethoprim-sulfamethoxazole test observations for which an identical training feature profile was associated with a conflicting phenotype label. Thus, identical feature representations were not invariably associated with identical susceptibility outcomes.

### 3.6 Performance on Nonoverlapping Test Isolates

The selected models were reevaluated after excluding test isolates with genomic feature profiles identical to those in their corresponding training datasets. No models were retrained. The remaining test observations therefore provided a direct sensitivity analysis of the original predictions.

**Table 2.** Model performance after exclusion of overlapping genomic feature profiles.

| **Antibiotic** | **Test n** | **Accuracy (%)** | **ROC-AUC** | **Resistant recall** |
|---|---|---|---|---|
| Amoxicillin-clavulanic acid | 22 | 86.36 | 0.915 | 0.929 |
| Ampicillin | 68 | 92.65 | 0.848 | 0.946 |
| Cefazolin | 28 | 67.86 | 0.795 | 0.533 |
| Cefotaxime | 20 | 85.00 | 1.000 | 0.700 |
| Cefoxitin | 8 | 100.00 | 1.000 | 1.000 |
| Ceftazidime | 43 | 83.72 | 0.908 | 0.222 |
| Ceftriaxone | 65 | 81.54 | 0.880 | 0.500 |
| Piperacillin-tazobactam | 22 | 90.91 | 1.000 | 0.500 |
| Trimethoprim-sulfamethoxazole | 51 | 90.20 | 0.901 | 0.925 |

**Figure 3.**
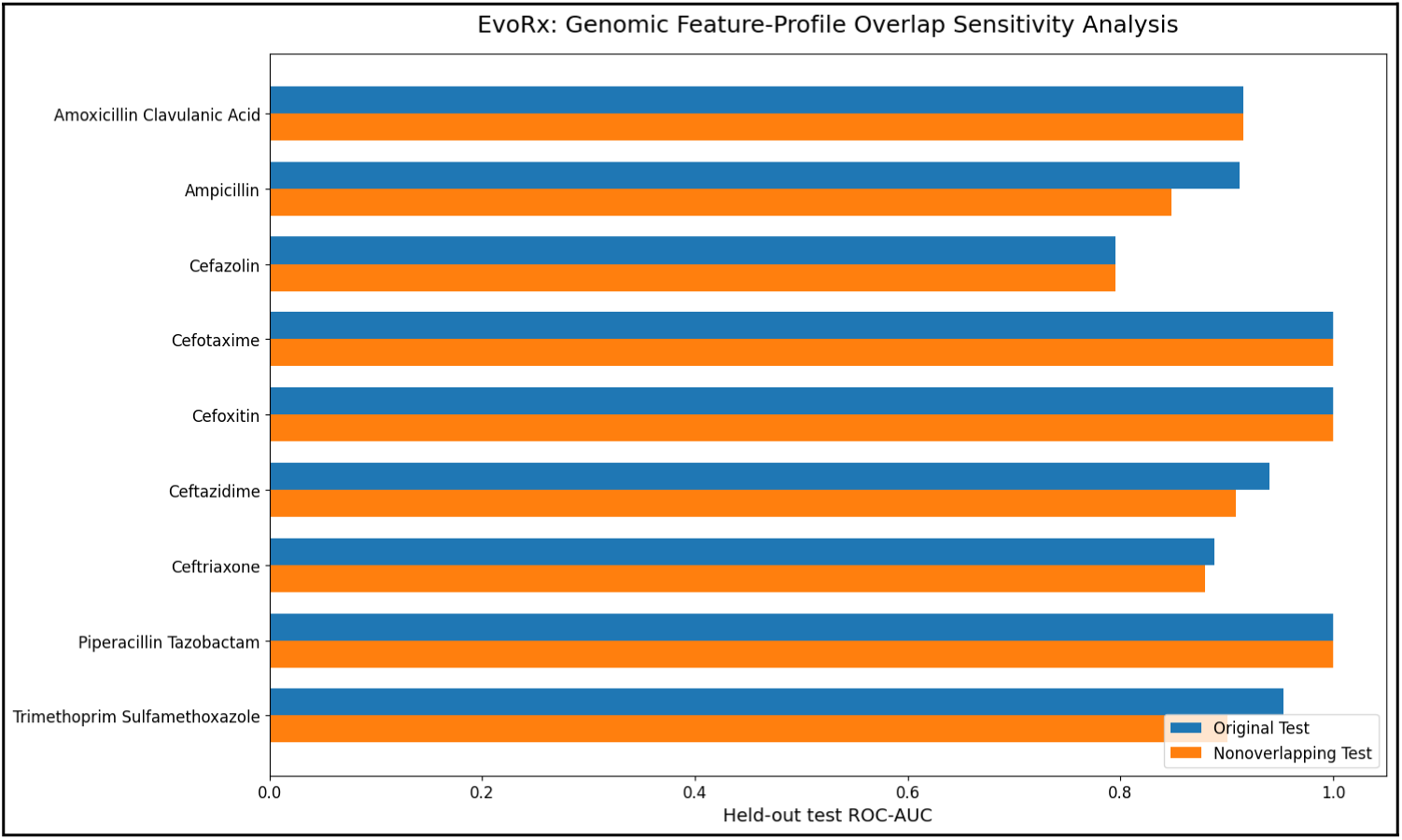
Comparison of held-out test ROC-AUC before and after exclusion of test isolates sharing identical 252-feature genomic profiles with training isolates. Previously fitted models were evaluated without retraining. Antibiotics without detected profile overlap retained unchanged evaluation subsets.

For ampicillin, exclusion of 15 overlapping observations reduced accuracy from 93.98% to 92.65% and ROC-AUC from 0.912 to 0.848. Resistant recall changed from 0.952 to 0.946.

For trimethoprim sulfamethoxazole, removal of 27 overlapping observations reduced accuracy from 93.59% to 90.20% and ROC-AUC from 0.954 to 0.901. Resistant recall declined from 0.945 to 0.925.

Ceftazidime accuracy decreased from 88.52% to 83.72%, while ROC-AUC declined from 0.940 to 0.908. Resistant recall remained 0.222. Ceftriaxone accuracy decreased from 83.78% to 81.54%, with ROC-AUC declining from 0.889 to 0.880.

Cefoxitin retained perfect classification of its eight remaining test observations. Performance for amoxicillin-clavulanic acid, cefazolin and cefotaxime was unchanged because no overlapping feature profiles had been identified.

### 3.7 Exploratory Cross-Publication Validation

Cross-publication evaluation comprised 18 eligible publication-held-out experiments across seven antibiotics. For each experiment, isolates from one source publication were evaluated using a model trained on isolates from other publications. Performance varied substantially between antibiotics and publication-defined subsets.

**Figure 4.**
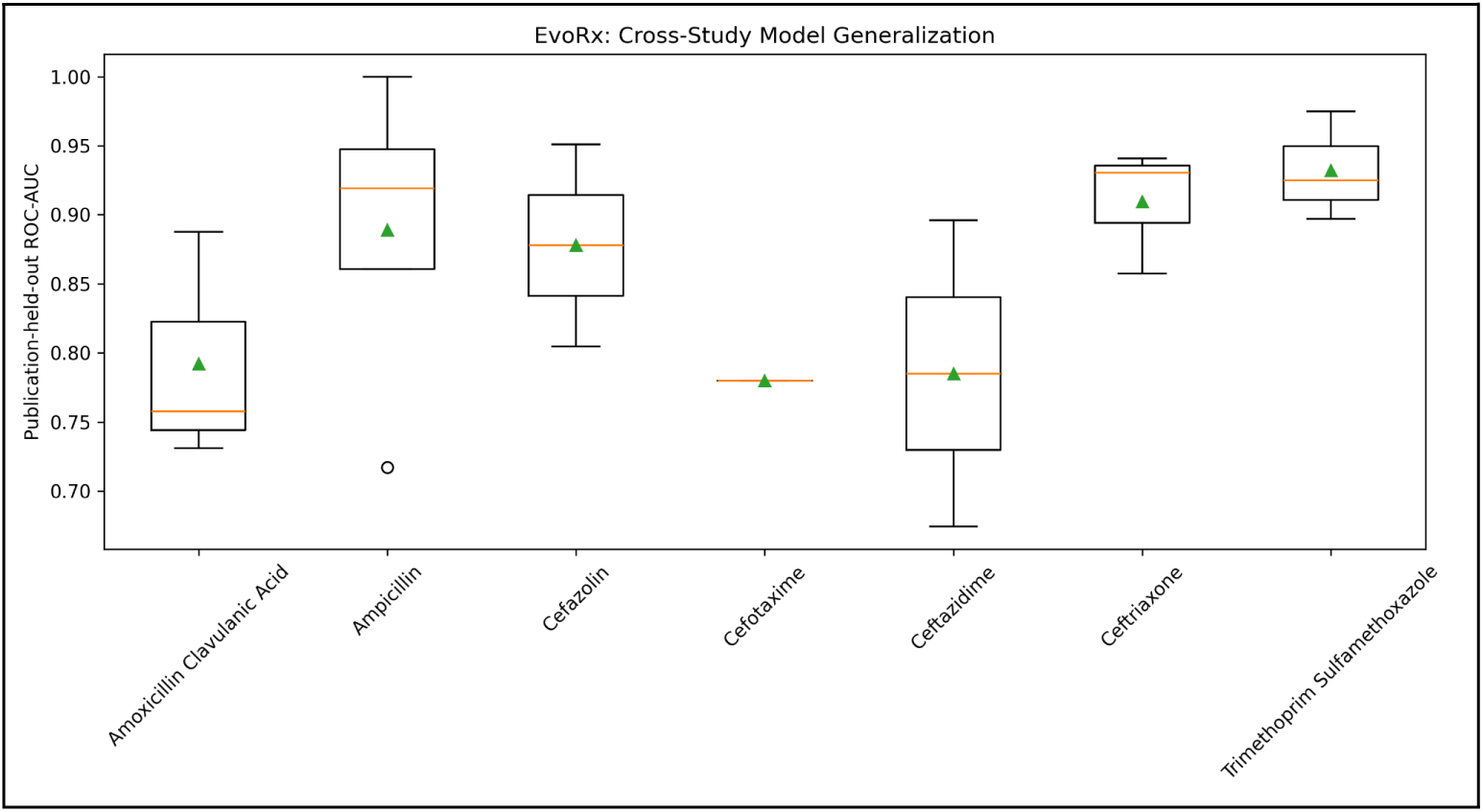
Distribution of ROC-AUC values across exploratory publication-held-out evaluations for seven antibiotics. Boxplots summarise performance across eligible publication-defined subsets. These evaluations were not fully independent at the lineage level, as shared MLST sequence types were identified across training and evaluation publications in 16 of 18 experiments.

Ampicillin was evaluated across four held-out publications, with accuracy ranging from 79.07% to 100.00% and ROC-AUC from 0.717 to 1.000. Resistant-isolate recall ranged from 0.654 to 1.000. For the largest ampicillin evaluation subset (n = 167), accuracy was 90.42%, balanced accuracy was 0.906, and ROC-AUC was 0.908.

Amoxicillin-clavulanic acid showed greater variation across three publication-held-out experiments. Accuracy ranged from 28.57% to 74.42%, while ROC-AUC ranged from 0.731 to 0.888. One evaluation subset containing 14 isolates yielded no correctly identified resistant isolates, despite a ROC-AUC of 0.888.

For ceftazidime, accuracy was 54.55% and 84.76% across two publication-held-out experiments. Resistant recall was zero in both, although ROC-AUC values were 0.896 and 0.674, respectively. Ceftriaxone accuracy ranged from 45.83% to 97.60% across three experiments. The remaining evaluations comprised two cefazolin experiments, one cefotaxime experiment and three trimethoprim-sulfamethoxazole experiments.

The accompanying lineage audit identified MLST sequence types shared between training and evaluation publications in 16 of the 18 experiments. No shared BioSample accessions or isolate-group identifiers were identified in these comparisons. Thus, publication-based separation did not consistently produce MLST-independent evaluation subsets.

### 3.8 Resistance-Associated Genomic Features

Associations between individual genomic determinants and phenotypic resistance were examined using gene prevalence and model-derived feature importance.

For ampicillin, blaTEM-1 was detected in 200 of 312 resistant isolates (64.10%), compared with 4 of 104 susceptible isolates (3.85%). Other beta-lactamase determinants were also more prevalent among resistant isolates. blaCTX-M-15 occurred in 12.50% of resistant isolates and none of the susceptible isolates, while blaCMY-2 was detected in 8.01% of resistant isolates and none of the susceptible isolates. blaOXA-1 was present in 11.54% and 0.96% of resistant and susceptible isolates, respectively.

For trimethoprim-sulfamethoxazole, sul2 was detected in 67.65% of resistant isolates and 13.16% of susceptible isolates. Corresponding prevalences were 51.10% and 7.89% for sul7, 22.43% and 2.63% for dfrA14, and 9.93% and 0.88% for dfrA8. These determinants also appeared among the strongest positive coefficients in the selected Logistic Regression model. The largest positive coefficients were associated with sul2 (2.426), sul1 (1.524), dfrA14 (1.234) and d/rA8 (1.161).

**Figure 5.**
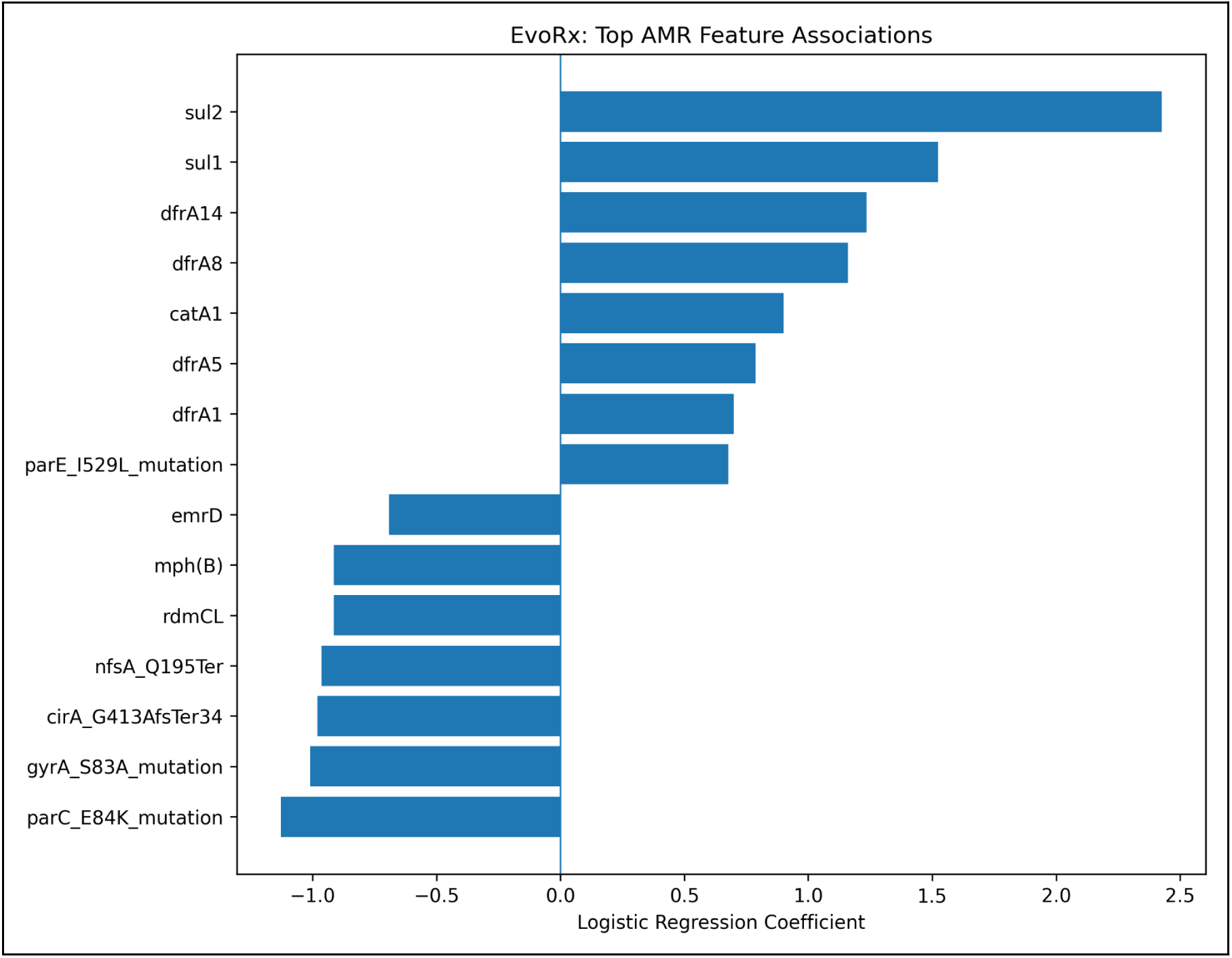
Logistic Regression coefficients for selected genomic features in the trimethoprim–sulfamethoxazole resistance model. Positive coefficients indicate associations with the resistant phenotype, whereas negative coefficients indicate associations with the susceptible phenotype, conditional on other model features. Coefficients do not establish causal effects.

Feature-importance profiles differed between antibiotics. In the Random Forest model for amoxicillin-clavulanic acid, blaCMY-2 had the highest impurity-based importance (0.089), followed by aph(3”)-lb (0.068) and aph(6)-Id (0.068). Several determinants associated with other antibiotic classes also received nonzero importance scores. These results describe predictive associations within the fitted models rather than independent evidence of causal resistance mechanisms.

### 3.9 Feature Redundancy and Model Robustness

Feature-redundancy analyses were conducted for ampicillin and trimethoprim-sulfamethoxazole by comparing the original 252-feature matrices with reduced representations.

For ampicillin, the feature count decreased from 252 to 144, a reduction of 42.86%. Validation accuracy remained 85.88%, balanced accuracy remained 0.905, and ROC-AUC remained 0.901. No change was observed in the recorded validation metrics.

For trimethoprim-sulfamethoxazole, the feature count decreased from 252 to 138, representing a 45.24% reduction. Validation accuracy remained 82.89%, and balanced accuracy remained 0.785. ROC-AUC increased slightly from 0.878 to 0.883, a difference of 0.0059.

**Table 3.**
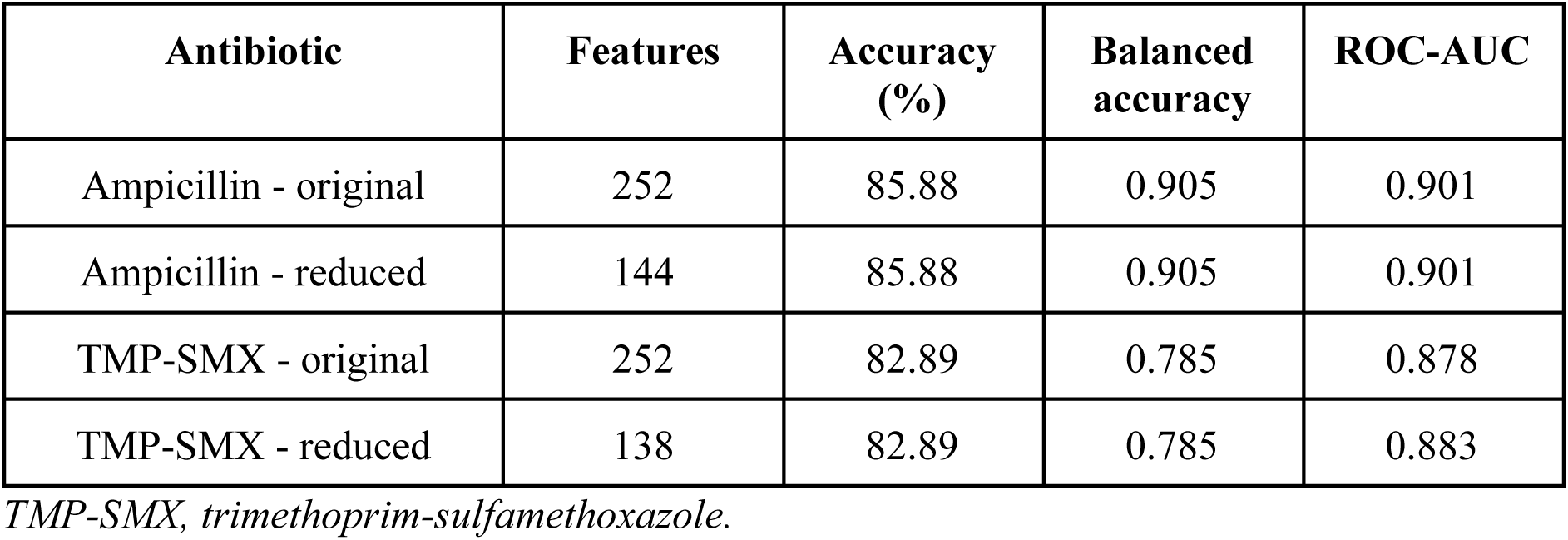
Validation performance before and after feature reduction.

These comparisons were performed on the validation datasets. The reduced feature representations were not subjected to separate independent test-set or cross-publication evaluations.

## 4. Discussion

### 4.1 Summary and Interpretation of Principal Findings

The EvoRx models demonstrated that resistance-associated genomic features can support the prediction of phenotypic antibiotic resistance in Escherichia coli, although performance differed considerably across antibiotics and evaluation conditions. Held-out accuracy exceeded 90% for ampicillin, cefoxitin, piperacillin-tazobactam and trimethoprim-sulfamethoxazole. However, differences between overall accuracy, ROC-AUC and resistant-isolate recall revealed substantial variation in the models’ ability to identify resistant infections.

The contrast was particularly evident for ceftazidime. Despite achieving 88.52% accuracy and a ROC-AUC of 0.940, the model identified only 22.22% of resistant isolates. Ceftriaxone and piperacillin-tazobactam similarly achieved relatively high overall accuracies while detecting only half of the resistant isolates. These findings demonstrate that strong aggregate classification metrics do not necessarily correspond to reliable resistance detection. ROC-AUC describes discrimination across possible thresholds, whereas recall reflects performance at the threshold used for classification. Their divergence is especially relevant when resistant isolates constitute a minority of the evaluation population.

The genomic overlap analysis provided a second distinction between apparent predictive performance and generalisability. All 74 test observations sharing identical 252-feature profiles with training isolates were classified correctly. Following their exclusion, ROC-AUC decreased from 0.912 to 0.848 for ampicillin and from 0.954 to 0.901 for trimethoprim-sulfamethoxazole. Although discrimination remained above chance, the reductions indicate that performance estimates were sensitive to the composition of the held-out datasets. Because the models were not retrained, these comparisons isolate changes in the evaluation population rather than changes in model fitting.

Cross-publication experiments further demonstrated variation in predictive performance, including cases of poor resistant-isolate recall. However, shared MLST sequence types occurred across training and evaluation publications in 16 of 18 experiments. Publication separation therefore did not consistently establish genetic independence. Taken together, the results support the predictive relevance of known genomic resistance determinants while distinguishing conventional held-out performance from evidence of generalisation to genetically unfamiliar bacterial populations.

### 4.2 Biological Significance of Resistance-Associated Features

The associations identified between genomic features and susceptibility phenotypes were broadly consistent with established mechanisms of antimicrobial resistance. For ampicillin, blaTEM-1 was detected in 64.10% of resistant isolates compared with 3.85% of susceptible isolates. This marked difference accords with the established contribution of TEM-family beta-lactamases to resistance against aminopenicillins [4]. Additional beta-lactamase genes, including blaCTX-M-15, blaCMY-2 and blaOXA-1, were more prevalent among resistant isolates, reflecting the diversity of enzymatic mechanisms represented in the dataset.

Nevertheless, the presence of a resistance determinant was not sufficient to account for every observed phenotype. Approximately 36% of ampicillin-resistant isolates lacked blaTEM-1, while a small proportion of susceptible isolates carried it. Other beta-lactamases, changes in permeability, regulatory variation and differences in gene functionality may contribute to these discrepancies. The analysis therefore supports the use of multiple genomic features rather than treating individual gene detections as universally definitive predictors.

For trimethoprim-sulfamethoxazole, the higher prevalence of sul1, sul2, dfrA14 and dfrA8 among resistant isolates was consistent with resistance mechanisms involving altered folate-pathway enzymes. These determinants also received positive coefficients in the selected Logistic Regression model. Their agreement with established biological mechanisms provides a degree of interpretability, although coefficient magnitude is influenced by feature correlations and model regularisation and cannot be interpreted as a direct measure of biological effect.

The Random Forest results also assigned importance to determinants not directly associated with the antibiotic being predicted. Such relationships may arise through co-occurrence of resistance genes on mobile genetic elements or through shared bacterial lineage backgrounds. Consequently, predictive importance does not establish that a feature independently causes resistance to the corresponding antibiotic. Distinguishing mechanistic determinants from correlated genomic markers remains necessary when interpreting feature-based resistance models.

### 4.3 Comparison with Existing Genomic AMR Prediction Studies

The findings are consistent with previous research demonstrating that bacterial genomic data can be used to predict antimicrobial susceptibility. Moradigaravand et al. investigated resistance prediction in E. coli using large-scale pan-genome information, establishing the relevance of genomic variation to machine-learning classification [10]. Pearcy et al. extended this approach by integrating machine learning with genome-scale metabolic modelling to investigate genetic determinants associated with antibiotic resistance [11]. EvoRx differs in its use of a restricted representation of 252 resistance-associated genes and mutations, rather than broader pan-genomic or metabolic features.

This distinction has methodological consequences. A curated resistance-feature representation provides a direct connection to established resistance biology and reduces the dimensionality of the genomic input. However, it may omit accessory genomic variation, regulatory changes and lineage-associated characteristics that contribute to susceptibility. The present results cannot establish whether the restricted representation performs better or worse than genome-wide approaches without a direct comparison using the same isolates, phenotypes and evaluation procedure.

Kim et al. identified data quality, model interpretability and clinical generalisation as important challenges in machine-learning-based AMR prediction [12]. The EvoRx results illustrate these concerns at the level of individual antibiotic outcomes. High overall accuracy did not consistently correspond to high resistant-isolate sensitivity, while the overlap analysis demonstrated that evaluation results depended partly on whether previously observed genomic feature profiles were represented in the test data.

These observations also align with broader methodological evidence that conventional cross-validation can yield misleading estimates when observations possess hierarchical, spatial or phylogenetic relationships [13]. In bacterial datasets, such dependence may arise from shared ancestry and the nonrandom distribution of resistance determinants. EvoRx contributes an antibiotic-specific examination of these effects within a curated genomic feature representation, rather than establishing a new class of resistance prediction methodology.

### 4.4 Potential Research Applications

The EvoRx framework provides a basis for investigating genomic predictors of antimicrobial resistance across multiple antibiotics. Its principal research application lies in evaluating how established resistance determinants contribute to phenotypic classification and whether these associations remain predictive across different isolate populations.

The use of a standardised binary feature representation also permits comparisons between antibiotics and modelling approaches. Such analyses could support the identification of resistance mechanisms that are consistently informative, as well as phenotypes for which existing genomic annotations provide insufficient predictive information. The observed differences in resistant-isolate recall suggest that antibiotic-specific evaluation is particularly important when assessing potential applications.

The computational workflow could also be adapted for genomic surveillance research, allowing newly sequenced isolates to be compared against previously characterised resistance profiles. However, EvoRx currently requires preprocessed genomic features and has not undergone prospective clinical validation. Its predictions should therefore be regarded as research outputs rather than substitutes for laboratory susceptibility testing.

### 4.5 Study Limitations

- Dataset size and unequal antibiotic coverage: Although the dataset comprised 446 E. coli genomes, susceptibility data varied across antibiotics. Small test subsets, particularly for cefoxitin (n = 9) and cefotaxime (n = 20), reduced the precision of performance estimates.
- Phenotypic heterogeneity: Susceptibility records originated from different laboratory sources and were not harmonised to a single testing protocol or breakpoint standard, potentially introducing variability in phenotype classifications. I
- Restricted genomic representation: The 252 binary features captured annotated resistance determinants but excluded potentially relevant information, including gene expression, regulatory variation and uncharacterised resistance mechanisms.
- Genomic overlap and limited phylogenetic independence: Seventy-four held-out test observations shared identical resistance-feature profiles with training isolates. Additionally, shared MLST sequence types occurred in 16 of 18 cross-publication experiments, limiting conclusions about generalisation to genetically independent populations.
- Exploratory validation: Cross-publication experiments did not employ fully nested model selection, and feature-reduction analyses lacked independent validation. Consequently, these findings provide preliminary evidence rather than definitive estimates of model robustness.
- Absence of prospective clinical validation: Models were evaluated retrospectively using existing genomic and phenotypic records. Their reliability on newly collected clinical isolates and previously unrepresented bacterial populations remains unestablished.

### 4.6 Future Research Directions

Further work should prioritise evaluation using larger, independently assembled datasets with harmonised susceptibility measurements and broader representation of resistant isolates. Validation based on genomic relatedness, including whole-genome phylogenetic clustering, would provide a more stringent assessment of generalisation than publication identifiers or MLST grouping alone.

Model development could also investigate approaches that address class imbalance and optimise resistance detection rather than overall accuracy. Threshold calibration, precision-recall analysis and antibiotic-specific sensitivity requirements would be particularly relevant for antibiotics such as ceftazidime, where strong ROC-AUC was accompanied by poor resistant-isolate recall.

Expanding the genomic representation to incorporate additional sequence variation and resistance-associated regulatory features may help identify determinants absent from the current annotation-based matrix. Comparisons with gene-presence baselines and broader pan-genomic models would clarify the contribution of machine learning beyond established resistance markers.

Finally, prospective evaluation on newly sequenced isolates, with independently measured phenotypic susceptibility, would be necessary to determine whether the observed predictive associations remain reliable outside the retrospective research dataset.

## 5. Conclusion

EvoRx demonstrates the potential of resistance-associated genomic features to predict phenotypic antibiotic resistance in Escherichia coli, while revealing substantial differences in predictive reliability across antibiotics. Using 252 binary genomic features derived from 446 genomes, the study evaluated Logistic Regression, Random Forest and XGBoost classifiers across nine antibiotics.

The strongest overall performance was observed for ampicillin and trimethoprim-sulfamethoxazole, which achieved held-out accuracies of 93.98% and 93.59%, respectively, alongside resistant-isolate recall exceeding 94%. However, high accuracy did not consistently indicate reliable resistance detection, as demonstrated by the substantially lower resistant recall for ceftazidime and ceftriaxone.

The genomic overlap analysis further established that conventional held-out performance was sensitive to shared resistance-feature profiles between training and test isolates. Cross-publication evaluation revealed additional variability, with residual lineage overlap limiting conclusions about generalisability.

These findings support genomic machine learning as a research approach for investigating antibiotic-specific resistance patterns, rather than an independently validated replacement for phenotypic susceptibility testing. Further evaluation using larger, phylogenetically independent datasets, harmonised susceptibility measurements and prospectively collected isolates is required to establish the robustness and potential clinical applicability of EvoRx.

## Declarations

### Data Availability

The study used publicly available Escherichia coli genomic assemblies and antimicrobial susceptibility records obtained through the Bacterial and Viral Bioinformatics Resource Centre (BV-BRC) and NCBI databases. The final analytical dataset comprised 446 genomes, 252 binary resistance-associated genomic features and 1,935 susceptibility observations across nine antibiotics.

The research archive contains the processed genomic feature matrices, antibiotic-specific phenotype datasets, dataset partition records and supplementary analysis outputs. Source genome accession identifiers and associated metadata are retained to support traceability and reproducibility.

### Code and Model Availability

The EvoRx computational workflow was implemented in Python using scikit-learn, XGBoost, pandas and NumPy. The research archive contains scripts and records for data preparation, model development, evaluation and supplementary analyses.

Trained classification models were preserved in Joblib format, together with feature-order records, validation comparisons, held-out test results and genomic overlap analyses. The computational workflow and saved model artefacts support reproduction of the reported analyses using the corresponding datasets and software dependencies.

### Author Contributions

A.A. conceived and designed the study, developed the research methodology, collected and curated genomic and antibiotic susceptibility data, performed data cleaning and quality control, and constructed the genomic matrix. A.A. developed and implemented the Python-based computational workflows, trained and evaluated different models. A.A. developed the figures and tables, interpreted the results, and prepared the original manuscript draft. E.N. supervised the research, provided methodological guidance and standard operating procedures, and offered feedback throughout the study. Both authors reviewed and approved the final manuscript.

## Acknowledgements

I would like to express my sincere gratitude to Dr Elamathi Natarajan for her mentorship, guidance, and encouragement throughout this research. I also acknowledge the researchers whose published studies and publicly available scientific resources provided a valuable foundation for this work.

