## Supplementary Information for "EvoRx: Machine Learning-Based Prediction of Antibiotic Resistance in Escherichia coli Using Genomic Features"

### SUPPLEMENTARY FIGURES

**Supplementary Figure S1. Validation and test performance of the Logistic Regression model**

Comparison of accuracy, balanced accuracy, and receiver operating characteristic area under the curve (ROC-AUC) for the Logistic Regression model across validation and held-out test datasets. Test performance exceeded validation performance for all three metrics. These results describe predictive performance under the evaluated data partitions and should be interpreted in the context of the genomic feature-profile overlap identified in the study.

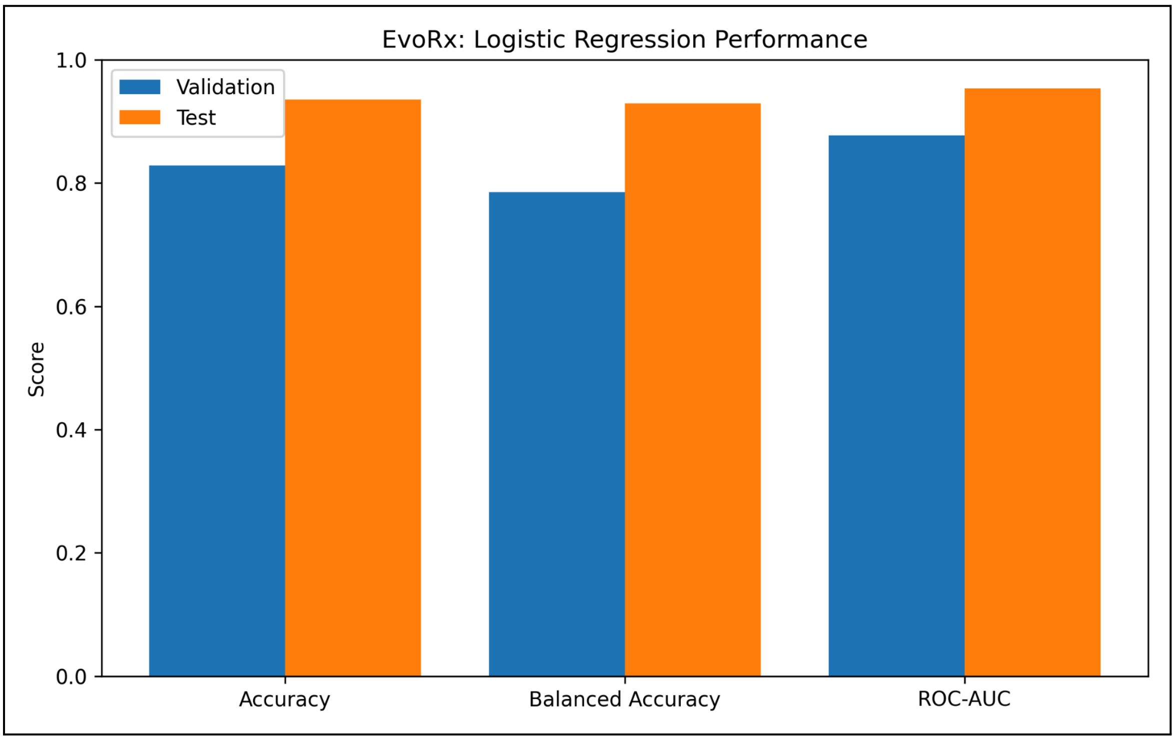

**Supplementary Figure S2. Publication-held-out evaluation of model performance**

ROC-AUC and balanced accuracy obtained from six publication-held-out evaluations, identified by their respective publication IDs. Predictive performance varied across publications, indicating differences in model generalisation between independently sourced datasets. These results are exploratory and should be interpreted alongside the number of observations, class distributions, and potential lineage overlap associated with each evaluation.

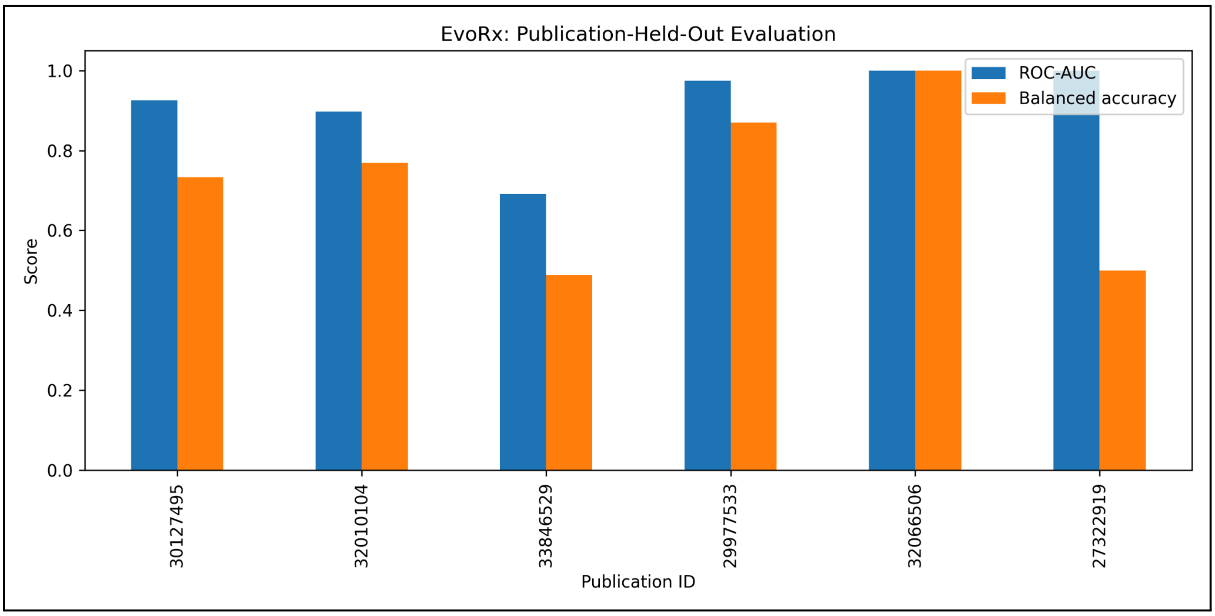

**Supplementary Figure S3. Sensitivity of model performance to genomic feature reduction**

Comparison of validation ROC-AUC before and after genomic feature reduction for ampicillin and trimethoprim-sulfamethoxazole. The original and reduced feature sets produced nearly identical performance for ampicillin, while a small improvement was observed following feature reduction for trimethoprim-sulfamethoxazole. These findings suggest that the evaluated models retained similar predictive performance after the removal of redundant genomic features.

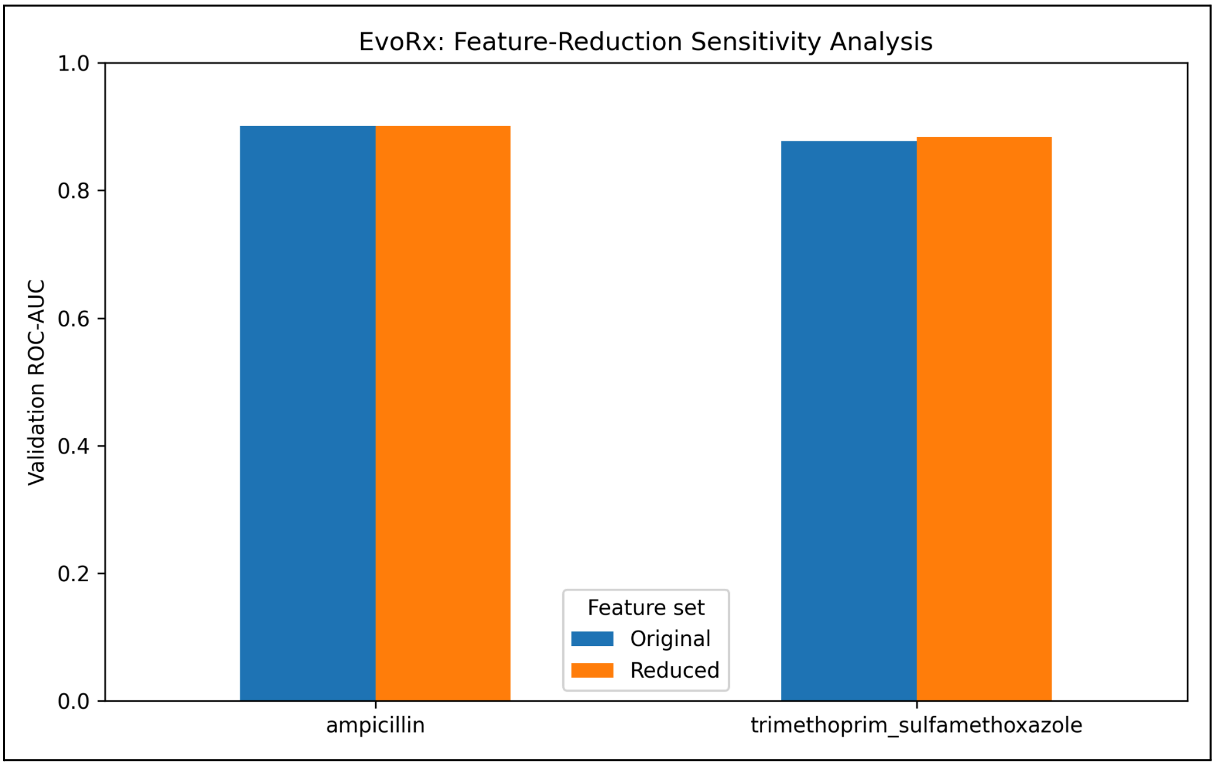

### SUPPLEMENTARY TABLES

**Supplementary Table S1. Dataset composition and class distribution across training, validation, and test partitions**

Distribution of resistant (R) and susceptible (S) observations across the training, validation, and held-out test partitions for each antibiotic included in the EvoRx analysis. The table reports the total number of observations and the corresponding resistant and susceptible class counts within each partition.

| Antibiotic | Split | Total | Resistant | Susceptible |
|---|---|---|---|---|
| amoxicillin_clavulanic_acid | train | 62 | 39 | 23 |
| amoxicillin_clavulanic_acid | validation | 21 | 13 | 8 |
| amoxicillin_clavulanic_acid | test | 22 | 14 | 8 |
| ampicillin | train | 248 | 187 | 61 |
| ampicillin | validation | 85 | 63 | 22 |
| ampicillin | test | 83 | 62 | 21 |
| cefazolin | train | 81 | 39 | 42 |
| cefazolin | validation | 26 | 13 | 13 |
| cefazolin | test | 28 | 15 | 13 |
| cefotaxime | train | 46 | 34 | 12 |
| cefotaxime | validation | 18 | 13 | 5 |
| cefotaxime | test | 20 | 10 | 10 |
| cefoxitin | train | 18 | 12 | 6 |
| cefoxitin | validation | 10 | 8 | 2 |
| cefoxitin | test | 9 | 7 | 2 |
| ceftazidime | train | 183 | 28 | 155 |
| ceftazidime | validation | 61 | 9 | 52 |
| ceftazidime | test | 61 | 9 | 52 |
| ceftriaxone | train | 198 | 43 | 155 |
| ceftriaxone | validation | 65 | 14 | 51 |
| ceftriaxone | test | 74 | 24 | 50 |
| piperacillin_tazobactam | train | 79 | 15 | 64 |

| | | | | |
|---|---|---|---|---|
| piperacillin_tazobactam | validation | 25 | 3 | 22 |
| piperacillin_tazobactam | test | 26 | 4 | 22 |
| trimethoprim_sulfamethoxazole | train | 232 | 163 | 69 |
| trimethoprim_sulfamethoxazole | validation | 76 | 54 | 22 |
| trimethoprim_sulfamethoxazole | test | 78 | 55 | 23 |

**Supplementary Table S2. Validation performance of machine-learning models across antibiotics**

Comparison of Logistic Regression, Random Forest, and XGBoost models on the validation datasets for the evaluated antibiotics. Performance is reported using accuracy, balanced accuracy, precision for the resistant class, recall for the resistant class, F1-score for the resistant class, and area under the receiver operating characteristic curve (ROC-AUC). The table also reports the total number of observations and the training and validation sample sizes. These results were used to compare candidate models before final held-out test evaluation.

| Antibiotic | Model | N_total | N_train | N_validation | Accuracy | Balanced_Accuracy | Precision_R | Recall_R | F1_R | ROC_AUC |
|---|---|---|---|---|---|---|---|---|---|---|
| amoxicillin_clavulanic_acid | Logistic Regression | 105 | 62 | 21 | 0.666667 | 0.610577 | 0.6875 | 0.846154 | 0.758621 | 0.706731 |
| amoxicillin_clavulanic_acid | Random Forest | 105 | 62 | 21 | 0.714286 | 0.625 | 0.684211 | 1 | 0.8125 | 0.764423 |
| amoxicillin_clavulanic_acid | XGBoost | 105 | 62 | 21 | 0.714286 | 0.625 | 0.684211 | 1 | 0.8125 | 0.658654 |
| ampicillin | Logistic Regression | 416 | 248 | 85 | 0.858824 | 0.904762 | 1 | 0.809524 | 0.894737 | 0.900794 |
| ampicillin | Random Forest | 416 | 248 | 85 | 0.811765 | 0.769481 | 0.885246 | 0.857143 | 0.870968 | 0.877345 |

| | | | | | | | | | | |
|---|---|---|---|---|---|---|---|---|---|---|
| ampicillin | XGBoost | 416 | 248 | 85 | 0.741176 | 0.766234 | 0.918367 | 0.714286 | 0.803571 | 0.822872 |
| cefazolin | Logistic Regression | 135 | 81 | 26 | 0.692308 | 0.692308 | 0.777778 | 0.538462 | 0.636364 | 0.857988 |
| cefazolin | Random Forest | 135 | 81 | 26 | 0.615385 | 0.615385 | 0.714286 | 0.384615 | 0.5 | 0.698225 |
| cefazolin | XGBoost | 135 | 81 | 26 | 0.653846 | 0.653846 | 0.7 | 0.538462 | 0.608696 | 0.686391 |
| cefotaxime | Logistic Regression | 84 | 46 | 18 | 0.611111 | 0.546154 | 0.75 | 0.692308 | 0.72 | 0.430769 |
| cefotaxime | Random Forest | 84 | 46 | 18 | 0.666667 | 0.461538 | 0.705882 | 0.923077 | 0.8 | 0.384615 |
| cefotaxime | XGBoost | 84 | 46 | 18 | 0.5 | 0.469231 | 0.7 | 0.538462 | 0.608696 | 0.369231 |
| cefoxitin | Logistic Regression | 37 | 18 | 10 | 0.7 | 0.4375 | 0.777778 | 0.875 | 0.823529 | 0.375 |
| cefoxitin | Random Forest | 37 | 18 | 10 | 0.7 | 0.4375 | 0.777778 | 0.875 | 0.823529 | 0.5 |
| cefoxitin | XGBoost | 37 | 18 | 10 | 0.7 | 0.4375 | 0.777778 | 0.875 | 0.823529 | 0.15625 |
| ceftazidime | Logistic Regression | 305 | 183 | 61 | 0.852459 | 0.59188 | 0.5 | 0.222222 | 0.307692 | 0.784188 |
| ceftazidime | Random Forest | 305 | 183 | 61 | 0.868852 | 0.555556 | 1 | 0.111111 | 0.2 | 0.855769 |

| | | | | | | | | | | |
|---|---|---|---|---|---|---|---|---|---|---|
| ceftazidime | XGBoost | 305 | 183 | 61 | 0.852459 | 0.54594 | 0.5 | 0.111111 | 0.181818 | 0.785256 |
| ceftriaxone | Logistic Regression | 337 | 198 | 65 | 0.830769 | 0.736695 | 0.615385 | 0.571429 | 0.592593 | 0.811625 |
| ceftriaxone | Random Forest | 337 | 198 | 65 | 0.876923 | 0.740196 | 0.875 | 0.5 | 0.636364 | 0.866246 |
| ceftriaxone | XGBoost | 337 | 198 | 65 | 0.907692 | 0.811625 | 0.9 | 0.642857 | 0.75 | 0.82423 |
| piperacillin_tazobactam | Logistic Regression | 130 | 79 | 25 | 0.88 | 0.5 | 0 | 0 | 0 | 1 |
| piperacillin_tazobactam | Random Forest | 130 | 79 | 25 | 0.88 | 0.5 | 0 | 0 | 0 | 0.984848 |
| piperacillin_tazobactam | XGBoost | 130 | 79 | 25 | 0.92 | 0.666667 | 1 | 0.333333 | 0.5 | 0.984848 |

**Supplementary Table S3. Held-out test performance of selected machine-learning models**

Performance of the selected machine-learning model for each antibiotic on its held-out test dataset. The table reports test sample size, resistant and susceptible class counts, accuracy, balanced accuracy, resistant-class precision, recall and F1-score, and ROC-AUC. Confusion matrix counts are also provided as true negatives (TN), false positives (FP), false negatives (FN), and true positives (TP). These results describe performance on the original held-out test partitions and should be interpreted alongside the genomic feature-profile overlap and sensitivity analyses reported in the study.

| Antibiotic | Selected_Model | Test_N | Test_R | Test_S | Accuracy | Balanced_Accuracy | Precision_R | Recall_R | F1_R | ROC_AUC | TN | FP | FN | TP |
|---|---|---|---|---|---|---|---|---|---|---|---|---|---|---|
| amoxicillin_clavulanic_acid | Random Forest | 22 | 14 | 8 | 0.863636 | 0.839286 | 0.866667 | 0.928571 | 0.896552 | 0.915179 | 6 | 2 | 1 | 13 |
| ampicillin | Logistic Regression | 83 | 62 | 21 | 0.939759 | 0.928187 | 0.967213 | 0.951613 | 0.95935 | 0.912442 | 19 | 2 | 3 | 59 |
| cefazolin | Logistic Regression | 28 | 15 | 13 | 0.678571 | 0.689744 | 0.8 | 0.533333 | 0.64 | 0.794872 | 11 | 2 | 7 | 8 |
| cefotaxime | Logistic Regression | 20 | 10 | 10 | 0.85 | 0.85 | 1 | 0.7 | 0.823529 | 1 | 10 | 0 | 3 | 7 |
| cefoxitin | Random Forest | 9 | 7 | 2 | 1 | 1 | 1 | 1 | 1 | 1 | 2 | 0 | 0 | 7 |
| ceftazidime | Random Forest | 61 | 9 | 52 | 0.885246 | 0.611111 | 1 | 0.222222 | 0.363636 | 0.940171 | 52 | 0 | 7 | 2 |

| | | | | | | | | | | | | | | |
|---|---|---|---|---|---|---|---|---|---|---|---|---|---|---|
| ceftriaxone | Random Forest | 74 | 24 | 50 | 0.837838 | 0.75 | 1 | 0.5 | 0.666667 | 0.88875 | 50 | 0 | 12 | 12 |
| piperacillin_tazobactam | Logistic Regression | 26 | 4 | 22 | 0.923077 | 0.75 | 1 | 0.5 | 0.666667 | 1 | 22 | 0 | 2 | 2 |

**Supplementary Table S4. Bootstrap 95% confidence intervals for held-out test performance**

Bootstrap-derived 95% confidence intervals for accuracy, balanced accuracy, and area under the receiver operating characteristic curve (ROC-AUC) across eight antibiotics evaluated on the held-out test datasets. The table reports the test sample size for each antibiotic and the lower and upper confidence limits for each performance metric. These intervals describe the uncertainty associated with the reported test performance estimates.

| Antibiotic | Test_N | Accuracy_CI_Lower | Accuracy_CI_Upper | Balanced_Accuracy_CI_Lower | Balanced_Accuracy_CI_Upper | ROC_AUC_CI_Lower | ROC_AUC_CI_Upper |
|---|---|---|---|---|---|---|---|
| amoxicillin_clavulanic_acid | 22 | 0.727273 | 1 | 0.651786 | 1 | 0.75 | 1 |
| ampicillin | 83 | 0.879518 | 0.987952 | 0.848694 | 0.991935 | 0.781855 | 1 |
| cefazolin | 28 | 0.5 | 0.821429 | 0.517949 | 0.833333 | 0.610256 | 0.943718 |
| cefotaxime | 20 | 0.7 | 1 | 0.7 | 1 | 1 | 1 |
| cefoxitin | 9 | 1 | 1 | 1 | 1 | 1 | 1 |
| ceftazidime | 61 | 0.852459 | 0.934426 | 0.5 | 0.777778 | 0.871741 | 0.987179 |
| ceftriaxone | 74 | 0.77027 | 0.905405 | 0.645833 | 0.854167 | 0.794125 | 0.968333 |
| piperacillin_tazobactam | 26 | 0.846154 | 1 | 0.5 | 1 | 1 | 1 |

**Supplementary Table S5. Genomic feature importance across antibiotic-specific prediction models**

Ranked genomic features identified by the antibiotic-specific machine-learning models. The table reports the antibiotic, feature rank, genomic feature identifier, importance value, and importance type. Feature importance values represent model-derived measures of predictive contribution and should not be interpreted as direct evidence of a causal relationship between a genomic feature and antibiotic resistance.

| Antibiotic | Rank | Feature | Importance | Importance_Type |
|---|---|---|---|---|
| amoxicillin_clavulanic_acid | 1 | blaCMY-2 | 0.089214 | Impurity importance |
| amoxicillin_clavulanic_acid | 2 | aph(3")-Ib | 0.067905 | Impurity importance |
| amoxicillin_clavulanic_acid | 3 | aph(6)-Id | 0.067735 | Impurity importance |
| amoxicillin_clavulanic_acid | 4 | sul2 | 0.042249 | Impurity importance |
| amoxicillin_clavulanic_acid | 5 | tet(B) | 0.036958 | Impurity importance |
| amoxicillin_clavulanic_acid | 6 | gyrA_D87N_mutation | 0.035103 | Impurity importance |
| amoxicillin_clavulanic_acid | 7 | parC_S80I_mutation | 0.034996 | Impurity importance |

| | | | | |
|---|---|---|---|---|
| amoxicillin_clavulanic_acid | 8 | gyrA_S83L_mutation | 0.034231 | Impurity importance |
| amoxicillin_clavulanic_acid | 9 | floR | 0.024673 | Impurity importance |
| amoxicillin_clavulanic_acid | 10 | mph(E) | 0.023599 | Impurity importance |
| amoxicillin_clavulanic_acid | 11 | blaCTX-M-32 | 0.020575 | Impurity importance |
| amoxicillin_clavulanic_acid | 12 | ble | 0.019694 | Impurity importance |
| amoxicillin_clavulanic_acid | 13 | mdtM | 0.018666 | Impurity importance |
| amoxicillin_clavulanic_acid | 14 | aadA2 | 0.018659 | Impurity importance |
| amoxicillin_clavulanic_acid | 15 | aph(3')-Ia | 0.018641 | Impurity importance |
| ampicillin | 1 | blaTEM-1 | 3.208766 | Logistic coefficient |
| ampicillin | 2 | blaCTX-M-1 | 2.435902 | Logistic coefficient |
| ampicillin | 3 | blaCMY-2 | 1.87121 | Logistic coefficient |
| ampicillin | 4 | blaCTX-M-15 | 1.411207 | Logistic coefficient |

| | | | | |
|---|---|---|---|---|
| ampicillin | 5 | blaOXA-1 | 1.149362 | Logistic coefficient |
| ampicillin | 6 | cyaA_S352T_mutation | 0.987475 | Logistic coefficient |
| ampicillin | 7 | parE_S458A_mutation | 0.936746 | Logistic coefficient |
| ampicillin | 8 | dfrA8 | 0.902667 | Logistic coefficient |
| ampicillin | 9 | parE_I355T_mutation | -0.83338 | Logistic coefficient |
| ampicillin | 10 | aac(3)-IId | 0.75471 | Logistic coefficient |
| ampicillin | 11 | tet(B) | 0.748835 | Logistic coefficient |
| ampicillin | 12 | aadA5 | -0.74385 | Logistic coefficient |
| ampicillin | 13 | qnrS1 | 0.695611 | Logistic coefficient |
| ampicillin | 14 | dfrA7 | 0.663507 | Logistic coefficient |
| ampicillin | 15 | mcr-1.1 | 0.646138 | Logistic coefficient |
| cefazolin | 1 | blaCTX-M-15 | 2.057132 | Logistic coefficient |
| cefazolin | 2 | blaCMY-2 | 1.009297 | Logistic coefficient |
| cefazolin | 3 | aac(3)-IIe | 0.894915 | Logistic coefficient |
| cefazolin | 4 | sul1 | 0.770975 | Logistic coefficient |
| cefazolin | 5 | mph(A) | 0.690707 | Logistic coefficient |
| cefazolin | 6 | mrx(A) | 0.690707 | Logistic coefficient |
| cefazolin | 7 | aac(6')-Ib3 | 0.591774 | Logistic coefficient |
| cefazolin | 8 | dfrA12 | 0.565618 | Logistic coefficient |
| cefazolin | 9 | sat2 | 0.557609 | Logistic coefficient |

| | | | | |
|---|---|---|---|---|
| cefazolin | 10 | aph(3'')-Ib | -0.54756 | Logistic coefficient |
| cefazolin | 11 | parC_E84A_mutation | 0.524969 | Logistic coefficient |
| cefazolin | 12 | blaKPC-3 | 0.524589 | Logistic coefficient |
| cefazolin | 13 | mdtM | -0.51736 | Logistic coefficient |
| cefazolin | 14 | nfsA_G126R_mutation | 0.479949 | Logistic coefficient |
| cefazolin | 15 | aac(3)-VIa | -0.47237 | Logistic coefficient |
| cefotaxime | 1 | mcr-1.1 | 0.827706 | Logistic coefficient |
| cefotaxime | 2 | tet(B) | 0.704517 | Logistic coefficient |
| cefotaxime | 3 | aadA1 | 0.70192 | Logistic coefficient |
| cefotaxime | 4 | armA | -0.68039 | Logistic coefficient |
| cefotaxime | 5 | parE_S458A_mutation | 0.64301 | Logistic coefficient |
| cefotaxime | 6 | aph(3')-Ia | -0.57434 | Logistic coefficient |
| cefotaxime | 7 | gyrA_D87N_mutation | 0.549739 | Logistic coefficient |
| cefotaxime | 8 | parC_S80I_mutation | 0.549739 | Logistic coefficient |
| cefotaxime | 9 | mdtM | -0.53431 | Logistic coefficient |
| cefotaxime | 10 | uhpT_E350Q_mutation | 0.520358 | Logistic coefficient |
| cefotaxime | 11 | dfrA12 | 0.506319 | Logistic coefficient |
| cefotaxime | 12 | aadA5 | -0.47103 | Logistic coefficient |
| cefotaxime | 13 | gyrA_S83L_mutation | 0.464091 | Logistic coefficient |

| | | | | |
|---|---|---|---|---|
| cefotaxime | 14 | ftsI_N337NYRIN_mutation | 0.422134 | Logistic coefficient |
| cefotaxime | 15 | blaCMY | 0.418413 | Logistic coefficient |
| cefoxitin | 1 | parC_S80I_mutation | 0.121559 | Impurity importance |
| cefoxitin | 2 | gyrA_D87N_mutation | 0.106267 | Impurity importance |
| cefoxitin | 3 | gyrA_S83L_mutation | 0.057892 | Impurity importance |
| cefoxitin | 4 | sul1 | 0.043053 | Impurity importance |
| cefoxitin | 5 | cyaA_S352T_mutation | 0.033447 | Impurity importance |
| cefoxitin | 6 | mdtM | 0.028019 | Impurity importance |
| cefoxitin | 7 | ble | 0.026954 | Impurity importance |
| cefoxitin | 8 | tet(B) | 0.026053 | Impurity importance |
| cefoxitin | 9 | blaCMY-2 | 0.022324 | Impurity importance |
| cefoxitin | 10 | blaCMY | 0.021755 | Impurity importance |
| cefoxitin | 11 | aph(3")-Ib | 0.021532 | Impurity importance |
| cefoxitin | 12 | parE_S458A_mutation | 0.020892 | Impurity importance |
| cefoxitin | 13 | dfrA8 | 0.018867 | Impurity importance |
| cefoxitin | 14 | aph(6)-Id | 0.018048 | Impurity importance |
| cefoxitin | 15 | blaTEM-1 | 0.017432 | Impurity importance |
| ceftazidime | 1 | blaCTX-M-15 | 0.062115 | Impurity importance |
| ceftazidime | 2 | blaOXA-1 | 0.051626 | Impurity importance |

| | | | | |
|---|---|---|---|---|
| ceftazidime | 3 | parC_S80I_mutation | 0.046906 | Impurity importance |
| ceftazidime | 4 | aac(6')-Ib-cr5 | 0.045982 | Impurity importance |
| ceftazidime | 5 | catB3 | 0.041469 | Impurity importance |
| ceftazidime | 6 | parC_E84V_mutation | 0.040354 | Impurity importance |
| ceftazidime | 7 | gyrA_D87N_mutation | 0.040184 | Impurity importance |
| ceftazidime | 8 | ptsI_V25I_mutation | 0.037844 | Impurity importance |
| ceftazidime | 9 | parE_I529L_mutation | 0.037627 | Impurity importance |
| ceftazidime | 10 | parE_S458A_mutation | 0.033695 | Impurity importance |
| ceftazidime | 11 | ftsI_N337NYRIN_mutation | 0.031863 | Impurity importance |
| ceftazidime | 12 | gyrA_S83L_mutation | 0.02762 | Impurity importance |
| ceftazidime | 13 | blaCMY-2 | 0.026984 | Impurity importance |
| ceftazidime | 14 | uhpT_E350Q_mutation | 0.024805 | Impurity importance |
| ceftazidime | 15 | aph(6)-Id | 0.024597 | Impurity importance |
| ceftriaxone | 1 | blaCTX-M-15 | 0.086282 | Impurity importance |
| ceftriaxone | 2 | gyrA_D87N_mutation | 0.056514 | Impurity importance |
| ceftriaxone | 3 | blaCMY-2 | 0.054536 | Impurity importance |
| ceftriaxone | 4 | parC_S80I_mutation | 0.051822 | Impurity importance |
| ceftriaxone | 5 | parE_S458A_mutation | 0.036108 | Impurity importance |
| ceftriaxone | 6 | aac(3)-IIe | 0.032457 | Impurity importance |

| | | | | |
|---|---|---|---|---|
| ceftriaxone | 7 | gyrA_S83L_mutation | 0.029385 | Impurity importance |
| ceftriaxone | 8 | emrD | 0.028469 | Impurity importance |
| ceftriaxone | 9 | sul2 | 0.027305 | Impurity importance |
| ceftriaxone | 10 | ftsI_N337NYRIN_mutation | 0.024493 | Impurity importance |
| ceftriaxone | 11 | ble | 0.023945 | Impurity importance |
| ceftriaxone | 12 | sul1 | 0.022185 | Impurity importance |
| ceftriaxone | 13 | catB3 | 0.021426 | Impurity importance |
| ceftriaxone | 14 | blaTEM-1 | 0.016973 | Impurity importance |
| ceftriaxone | 15 | aadA1 | 0.015844 | Impurity importance |
| piperacillin_tazobactam | 1 | ftsI_N337NYRIN_mutation | 1.288356 | Logistic coefficient |
| piperacillin_tazobactam | 2 | blaCMY-42 | 0.938486 | Logistic coefficient |
| piperacillin_tazobactam | 3 | blaKPC-3 | 0.869959 | Logistic coefficient |
| piperacillin_tazobactam | 4 | parE_S458A_mutation | 0.83257 | Logistic coefficient |
| piperacillin_tazobactam | 5 | ompF_N174insTer18 | 0.797867 | Logistic coefficient |
| piperacillin_tazobactam | 6 | blaCMY | 0.797867 | Logistic coefficient |
| piperacillin_tazobactam | 7 | ble | 0.752422 | Logistic coefficient |
| piperacillin_tazobactam | 8 | blaOXA-9 | 0.743852 | Logistic coefficient |
| piperacillin_tazobactam | 9 | aac(6')-Ib' | 0.725392 | Logistic coefficient |

| | | | | |
|---|---|---|---|---|
| piperacillin_tazobactam | 10 | dfrA14 | 0.64489 | Logistic coefficient |
| piperacillin_tazobactam | 11 | blaOXA-1 | 0.630255 | Logistic coefficient |
| piperacillin_tazobactam | 12 | cirA_S90YfsTer15_mutation | 0.58571 | Logistic coefficient |
| piperacillin_tazobactam | 13 | acrR_V29YfsTer44 | 0.58571 | Logistic coefficient |
| piperacillin_tazobactam | 14 | parE_I529L_mutation | 0.574801 | Logistic coefficient |
| piperacillin_tazobactam | 15 | parC_E84V_mutation | 0.574801 | Logistic coefficient |

**Supplementary Table S6. Sensitivity of model performance to genomic feature redundancy**

Comparison of model validation performance before and after genomic feature reduction for ampicillin and trimethoprim-sulfamethoxazole. The table reports the original and reduced feature counts alongside validation accuracy, balanced accuracy, and ROC-AUC. These analyses assess whether removing redundant genomic features changes predictive performance within the evaluated validation datasets.

| Antibiotic | Version | Features | Validation_Accuracy | Validation_Balanced_Accuracy | Validation_ROC_AUC |
|---|---|---|---|---|---|
| ampicillin | Original | 252 | 0.858824 | 0.904762 | 0.900794 |
| ampicillin | Reduced | 144 | 0.858824 | 0.904762 | 0.900794 |
| trimethoprim_sulfamethoxazole | Original | 252 | 0.828947 | 0.785354 | 0.877525 |
| trimethoprim_sulfamethoxazole | Reduced | 138 | 0.828947 | 0.785354 | 0.883418 |

**Supplementary Table S7. Publication-held-out evaluation of antibiotic resistance prediction models**

Performance of antibiotic-specific machine-learning models under publication-held-out evaluation. The table reports the antibiotic, held-out publication identifier, model type, training and test sample sizes, resistant and susceptible test observations, accuracy, balanced accuracy, resistant-class recall, and area under the receiver operating characteristic curve (ROC-AUC). These exploratory evaluations assess model performance when observations from a designated publication are excluded from training. Results should be interpreted in the context of differences in sample size, class distribution, and potential genomic lineage overlap between publications.

| Antibiotic | Held_Out_Publication | Model | Train_N | Test_N | Test_R | Test_S | Accuracy | Balanced_Accuracy | Recall_R | ROC_AUC |
|---|---|---|---|---|---|---|---|---|---|---|
| amoxicillin_clavulanic_acid | 29977533 | Random Forest | 60 | 24 | 13 | 11 | 0.5 | 0.51049 | 0.384615 | 0.730769 |
| amoxicillin_clavulanic_acid | 31053578 | Random Forest | 70 | 14 | 10 | 4 | 0.285714 | 0.5 | 0 | 0.8875 |
| amoxicillin_clavulanic_acid | 33846529 | Random Forest | 41 | 43 | 24 | 19 | 0.744186 | 0.732456 | 0.833333 | 0.757675 |
| ampicillin | 29977533 | Logistic Regression | 342 | 24 | 16 | 8 | 1 | 1 | 1 | 1 |
| ampicillin | 30127495 | Logistic Regression | 199 | 167 | 110 | 57 | 0.904192 | 0.90614 | 0.9 | 0.908453 |
| ampicillin | 32010104 | Logistic Regression | 276 | 90 | 73 | 17 | 0.866667 | 0.669621 | 0.986301 | 0.929895 |

| | | | | | | | | | | |
|---|---|---|---|---|---|---|---|---|---|---|
| ampicillin | 33846529 | Logistic Regression | 323 | 43 | 26 | 17 | 0.790698 | 0.826923 | 0.653846 | 0.717195 |
| cefazolin | 29977533 | Logistic Regression | 69 | 24 | 13 | 11 | 0.875 | 0.884615 | 0.769231 | 0.951049 |
| cefazolin | 32010104 | Logistic Regression | 26 | 67 | 14 | 53 | 0.835821 | 0.738544 | 0.571429 | 0.804582 |
| cefotaxime | 33846529 | Logistic Regression | 10 | 41 | 21 | 20 | 0.731707 | 0.728571 | 0.857143 | 0.779762 |
| ceftazidime | 29977533 | Random Forest | 250 | 22 | 10 | 12 | 0.545455 | 0.5 | 0 | 0.895833 |
| ceftazidime | 30127495 | Random Forest | 108 | 164 | 5 | 159 | 0.847561 | 0.437107 | 0 | 0.674214 |
| ceftriaxone | 29977533 | Random Forest | 259 | 24 | 13 | 11 | 0.458333 | 0.5 | 0 | 0.940559 |
| ceftriaxone | 30127495 | Random Forest | 116 | 167 | 5 | 162 | 0.976048 | 0.6 | 0.2 | 0.857407 |
| ceftriaxone | 32010104 | Random Forest | 191 | 92 | 13 | 79 | 0.891304 | 0.615385 | 0.230769 | 0.93038 |
| trimethoprim_sulfamethoxazole | 29977533 | Logistic Regression | 320 | 24 | 17 | 7 | 0.875 | 0.869748 | 0.882353 | 0.97479 |

| | | | | | | | | | | |
|---|---|---|---|---|---|---|---|---|---|---|
| trimethoprim_sulfamethoxazole | 30127495 | Logistic Regression | 178 | 166 | 108 | 58 | 0.662651 | 0.732759 | 0.5 | 0.924808 |
| trimethoprim_sulfamethoxazole | 32010104 | Logistic Regression | 254 | 90 | 58 | 32 | 0.811111 | 0.769397 | 0.913793 | 0.896821 |

**Supplementary Table S8. Sensitivity of held-out test performance to overlapping genomic feature profiles**

Comparison of original held-out test performance with performance calculated after excluding test observations whose genomic feature profiles matched those present in the training dataset. The table reports original and nonoverlapping sample sizes, the number and percentage of excluded observations, and changes in accuracy, balanced accuracy, resistant-class recall, and ROC-AUC for each antibiotic. These analyses assess the sensitivity of reported predictive performance to genomic feature-profile overlap between training and test partitions.

| Antibiotic | Original_N | Nonoverlap_N | Excluded_N | Excluded_Percent | Original_Accuracy | Nonoverlap_Accuracy | Change_Accuracy | Original_Balanced_Accuracy | Nonoverlap_Balanced_Accuracy | Change_Balanced_Accuracy | Original_Recall_R | Nonoverlap_Recall_R | Change_Recall_R | Original_ROC_AUC | Nonoverlap_ROC_AUC | Change_ROC_AUC | Interpretation |
|---|---|---|---|---|---|---|---|---|---|---|---|---|---|---|---|---|---|
| amoxicillin_clavulanic_acid | 22 | 22 | 0 | 0 | 0.863636 | 0.863636 | 0 | 0.839286 | 0.839286 | 0 | 0.928571 | 0.928571 | 0 | 0.915179 | 0.915179 | 0 | No overlapping profiles |
| ampicillin | 83 | 68 | 15 | 18.07229 | 0.939759 | 0.926471 | -0.01329 | 0.928187 | 0.889881 | -0.03831 | 0.951613 | 0.946429 | -0.00518 | 0.912442 | 0.848214 | -0.06423 | Substantial sensitivity |
| cefazolin | 28 | 28 | 0 | 0 | 0.678571 | 0.678571 | 0 | 0.689744 | 0.689744 | 0 | 0.533333 | 0.533333 | 0 | 0.794872 | 0.794872 | 0 | No overlapping profiles |

| | | | | | | | | | | | | | | | | | |
|---|---|---|---|---|---|---|---|---|---|---|---|---|---|---|---|---|---|
| cefotaxime | 20 | 20 | 0 | 0 | 0.85 | 0.85 | 0 | 0.85 | 0.85 | 0 | 0.7 | 0.7 | 0 | 1 | 1 | 0 | No overlapping profiles |
| cefoxitin | 9 | 8 | 1 | 11.11111 | 1 | 1 | 0 | 1 | 1 | 0 | 1 | 1 | 0 | 1 | 1 | 0 | Limited observed AUC change |
| ceftazidime | 61 | 43 | 18 | 29.5082 | 0.885246 | 0.837209 | -0.04804 | 0.611111 | 0.611111 | 0 | 0.222222 | 0.222222 | 0 | 0.940171 | 0.908497 | -0.03167 | Limited observed AUC change |
| ceftriaxone | 74 | 65 | 9 | 12.16216 | 0.837838 | 0.815385 | -0.02245 | 0.75 | 0.75 | 0 | 0.5 | 0.5 | 0 | 0.88875 | 0.879573 | -0.00918 | Limited observed AUC change |
| piperacillin_tazobactam | 26 | 22 | 4 | 15.38462 | 0.923077 | 0.909091 | -0.01399 | 0.75 | 0.75 | 0 | 0.5 | 0.5 | 0 | 1 | 1 | 0 | Limited observed AUC change |
| trimethoprim_sulfamethoxazole | 78 | 51 | 27 | 34.61538 | 0.935897 | 0.901961 | -0.03394 | 0.929249 | 0.871591 | -0.05766 | 0.945455 | 0.925 | -0.02045 | 0.953755 | 0.901136 | -0.05262 | Substantial sensitivity |

**Supplementary Table S9. Genomic profile and lineage overlap audit of held-out test observations**

Detailed audit of held-out test observations with genomic feature profiles identical to those of at least one training observation. For each observation, the table reports the antibiotic, test genome accession, susceptibility classification, number of matching training isolates, and whether matching observations share multilocus sequence type (MLST), publication, BioSample accession, or isolate-group identifier. These comparisons characterise the relationship between overlapping genomic profiles and available isolate metadata. Identical feature profiles do not necessarily establish that the underlying genomes or bacterial isolates are identical.

| Antibiotic | Test_Accession | Test_Target | Matching_Train_Isolates | Shared_mlst | Shared_publication | Shared_biosample_accession | Shared_isolate_group_id |
|---|---|---|---|---|---|---|---|
| ampicillin | GCA_009650035.1 | S | 9 | FALSE | FALSE | FALSE | FALSE |
| ampicillin | GCA_004283435.1 | R | 1 | FALSE | TRUE | FALSE | FALSE |
| ampicillin | GCA_900490685.1 | S | 9 | FALSE | FALSE | FALSE | FALSE |
| ampicillin | GCA_900406605.1 | S | 9 | FALSE | FALSE | FALSE | FALSE |
| ampicillin | GCA_900406625.1 | S | 9 | FALSE | FALSE | FALSE | FALSE |
| ampicillin | GCA_004284515.1 | R | 1 | FALSE | TRUE | FALSE | FALSE |
| ampicillin | GCA_001286605.1 | S | 1 | FALSE | FALSE | FALSE | FALSE |
| ampicillin | GCA_001286405.1 | S | 4 | FALSE | TRUE | FALSE | FALSE |
| ampicillin | GCA_001285805.1 | R | 1 | FALSE | TRUE | FALSE | FALSE |
| ampicillin | GCA_001284805.1 | R | 1 | FALSE | TRUE | FALSE | FALSE |
| ampicillin | GCA_001283185.1 | S | 9 | FALSE | TRUE | FALSE | FALSE |

| | | | | | | | |
|---|---|---|---|---|---|---|---|
| ampicillin | GCA_001284545.1 | S | 9 | FALSE | TRUE | FALSE | FALSE |
| ampicillin | GCA_001284485.1 | S | 9 | FALSE | TRUE | FALSE | FALSE |
| ampicillin | GCA_001283705.1 | R | 1 | FALSE | TRUE | FALSE | FALSE |
| ampicillin | GCA_001286205.1 | R | 1 | FALSE | TRUE | FALSE | FALSE |
| cefoxitin | GCA_009650035.1 | S | 1 | FALSE | FALSE | FALSE | FALSE |
| ceftazidime | GCA_001286125.1 | S | 9 | FALSE | TRUE | FALSE | FALSE |
| ceftazidime | GCA_001286105.1 | S | 3 | FALSE | TRUE | FALSE | FALSE |
| ceftazidime | GCA_001286045.1 | S | 3 | FALSE | TRUE | FALSE | FALSE |
| ceftazidime | GCA_001286605.1 | S | 1 | FALSE | FALSE | FALSE | FALSE |
| ceftazidime | GCA_001286585.1 | S | 3 | FALSE | TRUE | FALSE | FALSE |
| ceftazidime | GCA_001285325.1 | S | 5 | FALSE | TRUE | FALSE | FALSE |
| ceftazidime | GCA_001285225.1 | S | 5 | FALSE | TRUE | FALSE | FALSE |
| ceftazidime | GCA_001277475.1 | S | 5 | FALSE | TRUE | FALSE | FALSE |
| ceftazidime | GCA_001277395.1 | S | 5 | FALSE | TRUE | FALSE | FALSE |
| ceftazidime | GCA_001285185.1 | S | 9 | FALSE | TRUE | FALSE | FALSE |
| ceftazidime | GCA_001283645.1 | S | 9 | FALSE | TRUE | FALSE | FALSE |
| ceftazidime | GCA_001283425.1 | S | 3 | FALSE | TRUE | FALSE | FALSE |
| ceftazidime | GCA_001283385.1 | S | 3 | FALSE | TRUE | FALSE | FALSE |
| ceftazidime | GCA_001284605.1 | S | 9 | FALSE | TRUE | FALSE | FALSE |
| ceftazidime | GCA_001283965.1 | S | 3 | FALSE | TRUE | FALSE | FALSE |

| | | | | | | | |
|---|---|---|---|---|---|---|---|
| ceftazidime | GCA_001283805.1 | S | 3 | FALSE | TRUE | FALSE | FALSE |
| ceftazidime | GCA_001284345.1 | S | 3 | FALSE | TRUE | FALSE | FALSE |
| ceftazidime | GCA_001284205.1 | S | 5 | FALSE | TRUE | FALSE | FALSE |
| ceftriaxone | GCA_004283865.1 | S | 16 | FALSE | FALSE | FALSE | FALSE |
| ceftriaxone | GCA_900406605.1 | S | 1 | FALSE | FALSE | FALSE | FALSE |
| ceftriaxone | GCA_900490565.1 | S | 1 | FALSE | TRUE | FALSE | FALSE |
| ceftriaxone | GCA_900406625.1 | S | 1 | FALSE | FALSE | FALSE | FALSE |
| ceftriaxone | GCA_004284005.1 | S | 16 | FALSE | FALSE | FALSE | FALSE |
| ceftriaxone | GCA_001285005.1 | S | 16 | FALSE | TRUE | FALSE | FALSE |
| ceftriaxone | GCA_001284565.1 | S | 4 | FALSE | TRUE | FALSE | FALSE |
| ceftriaxone | GCA_001283945.1 | S | 4 | FALSE | TRUE | FALSE | FALSE |
| ceftriaxone | GCA_001283785.1 | S | 4 | FALSE | TRUE | FALSE | FALSE |
| piperacillin_tazobactam | GCA_009650035.1 | S | 1 | FALSE | FALSE | FALSE | FALSE |
| piperacillin_tazobactam | GCA_004283865.1 | S | 1 | FALSE | FALSE | FALSE | FALSE |
| piperacillin_tazobactam | GCA_004284005.1 | S | 1 | FALSE | FALSE | FALSE | FALSE |
| piperacillin_tazobactam | GCA_004283725.1 | S | 1 | FALSE | TRUE | FALSE | FALSE |
| trimethoprim_sulfamethoxazole | GCA_004283865.1 | S | 11 | FALSE | FALSE | FALSE | FALSE |
| trimethoprim_sulfamethoxazole | GCA_900490675.1 | S | 1 | FALSE | FALSE | FALSE | FALSE |

| | | | | | | | |
|---|---|---|---|---|---|---|---|
| trimethoprim_sulfamethoxazole | GCA_004284005.1 | S | 11 | FALSE | FALSE | FALSE | FALSE |
| trimethoprim_sulfamethoxazole | GCA_001286185.1 | R | 2 | FALSE | TRUE | FALSE | FALSE |
| trimethoprim_sulfamethoxazole | GCA_001286125.1 | S | 11 | FALSE | TRUE | FALSE | FALSE |
| trimethoprim_sulfamethoxazole | GCA_001286465.1 | S | 1 | FALSE | TRUE | FALSE | FALSE |
| trimethoprim_sulfamethoxazole | GCA_001286365.1 | R | 2 | FALSE | TRUE | FALSE | FALSE |
| trimethoprim_sulfamethoxazole | GCA_001285685.1 | R | 5 | FALSE | TRUE | FALSE | FALSE |
| trimethoprim_sulfamethoxazole | GCA_001286645.1 | R | 5 | FALSE | TRUE | FALSE | FALSE |
| trimethoprim_sulfamethoxazole | GCA_001285305.1 | R | 5 | FALSE | TRUE | FALSE | FALSE |
| trimethoprim_sulfamethoxazole | GCA_001285485.1 | R | 4 | FALSE | TRUE | FALSE | FALSE |
| trimethoprim_sulfamethoxazole | GCA_001277455.1 | S | 11 | FALSE | TRUE | FALSE | FALSE |

| | | | | | | | |
|---|---|---|---|---|---|---|---|
| trimethoprim_sulfamethoxazole | GCA_001277675.1 | S | 11 | FALSE | TRUE | FALSE | FALSE |
| trimethoprim_sulfamethoxazole | GCA_001277635.1 | R | 2 | FALSE | TRUE | FALSE | FALSE |
| trimethoprim_sulfamethoxazole | GCA_001277615.1 | R | 5 | FALSE | TRUE | FALSE | FALSE |
| trimethoprim_sulfamethoxazole | GCA_004283725.1 | S | 1 | FALSE | TRUE | FALSE | FALSE |
| trimethoprim_sulfamethoxazole | GCA_001285185.1 | S | 11 | FALSE | TRUE | FALSE | FALSE |
| trimethoprim_sulfamethoxazole | GCA_001284945.1 | R | 5 | FALSE | TRUE | FALSE | FALSE |
| trimethoprim_sulfamethoxazole | GCA_001283545.1 | R | 2 | FALSE | TRUE | FALSE | FALSE |
| trimethoprim_sulfamethoxazole | GCA_001283485.1 | R | 2 | FALSE | TRUE | FALSE | FALSE |
| trimethoprim_sulfamethoxazole | GCA_001284565.1 | R | 4 | FALSE | TRUE | FALSE | FALSE |
| trimethoprim_sulfamethoxazole | GCA_001284485.1 | S | 11 | FALSE | TRUE | FALSE | FALSE |

| | | | | | | | |
|---|---|---|---|---|---|---|---|
| trimethoprim_sulfamethoxazole | GCA_001283945.1 | R | 4 | FALSE | TRUE | FALSE | FALSE |
| trimethoprim_sulfamethoxazole | GCA_001283925.1 | S | 2 | FALSE | TRUE | FALSE | FALSE |
| trimethoprim_sulfamethoxazole | GCA_001283785.1 | R | 4 | FALSE | TRUE | FALSE | FALSE |
| trimethoprim_sulfamethoxazole | GCA_001283725.1 | S | 11 | FALSE | TRUE | FALSE | FALSE |
| trimethoprim_sulfamethoxazole | GCA_001284065.1 | R | 2 | FALSE | TRUE | FALSE | FALSE |

**Supplementary Table S10. Comprehensive held-out test performance and 95% confidence intervals across nine antibiotics**

Summary of the selected machine-learning model and held-out test performance for each of the nine antibiotics evaluated in EvoRx. The table reports test sample sizes, resistant (R) and susceptible (S) class distributions, accuracy, balanced accuracy, resistant-class precision, recall, F1-score, and area under the receiver operating characteristic curve (ROC-AUC). Confusion matrix counts are provided as true negatives (TN), false positives (FP), false negatives (FN), and true positives (TP). Lower and upper 95% confidence interval bounds are reported for accuracy, balanced accuracy, and ROC-AUC. These results describe predictive performance on the original held-out test partitions and should be interpreted alongside the genomic feature-profile overlap and sensitivity analyses.

| Antibiotic | Selected_Model | Test_N | Test_R | Test_S | Accuracy | Balanced_Accuracy | Precision_R | Recall_R | F1_R | ROC_AUC | TN | FP | FN | TP | Accuracy_CI_Lower | Accuracy_CI_Upper | Balanced_Accuracy_CI_Lower | Balanced_Accuracy_CI_Upper | ROC_AUC_CI_Lower | ROC_AUC_CI_Upper |
|---|---|---|---|---|---|---|---|---|---|---|---|---|---|---|---|---|---|---|---|---|
| amoxicillin_clavulanic_acid | Random Forest | 22 | 14 | 8 | 0.863636 | 0.839286 | 0.866667 | 0.928571 | 0.896552 | 0.915179 | 6 | 2 | 1 | 13 | 0.727273 | 1 | 0.651786 | 1 | 0.75 | 1 |
| ampicillin | Logistic Regression | 83 | 62 | 21 | 0.939759 | 0.928187 | 0.967213 | 0.951613 | 0.959135 | 0.912442 | 19 | 2 | 3 | 59 | 0.879518 | 0.987952 | 0.848694 | 0.991935 | 0.781855 | 1 |

| | | | | | | | | | | | | | | | | | | | | |
|---|---|---|---|---|---|---|---|---|---|---|---|---|---|---|---|---|---|---|---|---|
| cefazolin | Logistic Regression | 28 | 15 | 13 | 0.678571 | 0.689744 | 0.8 | 0.533333 | 0.64 | 0.794872 | 11 | 2 | 7 | 8 | 0.5 | 0.821429 | 0.517949 | 0.833333 | 0.610256 | 0.943718 |
| cefotaxime | Logistic Regression | 20 | 10 | 10 | 0.85 | 0.85 | 1 | 0.7 | 0.823529 | 1 | 10 | 0 | 3 | 7 | 0.7 | 1 | 0.7 | 1 | 1 | 1 |
| cefoxitin | Random Forest | 9 | 7 | 2 | 1 | 1 | 1 | 1 | 1 | 1 | 2 | 0 | 0 | 7 | 1 | 1 | 1 | 1 | 1 | 1 |
| ceftazidime | Random Forest | 61 | 9 | 52 | 0.885246 | 0.611111 | 1 | 0.222222 | 0.363636 | 0.940171 | 52 | 0 | 7 | 2 | 0.852459 | 0.934426 | 0.5 | 0.777778 | 0.871741 | 0.987179 |
| ceftriaxone | Random Forest | 74 | 24 | 50 | 0.837838 | 0.75 | 1 | 0.5 | 0.666667 | 0.88875 | 50 | 0 | 12 | 12 | 0.77027 | 0.905405 | 0.645833 | 0.854167 | 0.794125 | 0.968333 |
| piperacillin_tazobactam | Logistic Regression | 26 | 4 | 22 | 0.923077 | 0.75 | 1 | 0.5 | 0.666667 | 1 | 22 | 0 | 2 | 2 | 0.846154 | 1 | 0.5 | 1 | 1 | 1 |
| trimethoprim_sulfamethoxazole | Logistic Regression | 78 | 55 | 23 | 0.935897 | 0.929249 | 0.962963 | 0.945455 | 0.954128 | 0.953755 | 21 | 2 | 3 | 52 | 0.871795 | 0.987179 | 0.854941 | 0.981818 | 0.869951 | 1 |
